# In Silico Discovery of Multi-Target Natural Ligands and Efficient siRNA Design for Overcoming Drug Resistance in Breast Cancer via Local Therapy

**DOI:** 10.1101/2024.11.10.622866

**Authors:** Seyed Mohammad Javad Hashemi, Ali Barzegar, Javad Akhtari, Amir Mellati

**Author notes:** **Corresponding Author:** Seyed Mohammad Javad Hashemi, Molecular Cell Biology Laboratory, Department of Basic Sciences, Sari Agricultural Sciences and Natural Resources University, Sari, Iran. Tell: 0098 9011098079.

## Abstract

In this study, we designed an efficient siRNA for PKMYT1 gene knockdown, and evaluated the binding affinities of different natural ligands to crucial proteins involved in breast cancer. Designed siRNA showed strong binding affinity and minimal off-target effects. Molecular docking studies identified new ligands as antagonists with high binding affinities for aromatase, estrogen receptor alpha, HER2, and PARP10, as well as agonists for MT2 and STING. The natural ligand SCHEMBL7562664 was introduced as a golden ligand due to its high affinity among multiple targets and lack of cytotoxic and mutagenic effects. Natural small molecules identified in this research, due to their multi-target characteristics, provided a solution to overcome the problem of drug resistance in cancer cells. Furthermore, the proposed three dimensional scaffold design for local breast cancer therapy offers a promising approach to increase the delivery and efficacy of these natural small molecules, reduce systemic side effects, and improve treatment outcomes. In this study, new ligands with significant binding affinities and favorable pharmacokinetic properties were identified, which paves the way for further research in targeted therapy of breast cancer.

## 1. Introduction

Breast cancer is the most common type of cancer in women, and according to the estimate of the American Cancer Society, 35 million people worldwide will be affected by it by 2050 (1). This disease has a multifactorial origin; environmental, and genetic factors play a role in its development (2,3).

If breast cancer is diagnosed in the early stages, with standard treatments such as chemotherapy, hormone, and radiation therapy, patients will have a higher chance of being cured(4,5).

Small molecules of natural or chemical origin can be used to modulate cell signaling pathways to target cancer. To reduce the side effects and biological toxicity caused by chemical agonists and antagonists, using natural types is a priority (6,7,8).

In silico studies as a computational method has provided the possibility of virtual screening to find effective drugs from a wide range of compounds. With bioinformatics modeling, it is possible to evaluate the interaction of compounds with specific cell receptors, as well as evaluate the safety and efficacy of new compounds. Such evaluations significantly reduce the time to discover new compounds for targeted cancer treatment, as well as the laboratory costs.

siRNA is a short and double-stranded RNA molecule that participates in the regulation of gene expression by destroying specific mRNA and, as a result, preventing the production of the corresponding protein. Their discovery in 1998 created a valuable tool in cancer treatment research, by targeting and reducing the expression of oncogenes, thus inhibiting cancer progression (9,10).

It is very beneficial to use local cancer treatment to increase the efficiency of treatment and minimize systemic side effects. In this method, three-dimensional (3D) scaffolds can be used for the gradual and stable release of drugs on-site. Previous studies have shown that using 3D-printed scaffolds carrying pharmaceutical compounds increased the success in reducing tumor size and reducing the toxic effects of drugs in the body (11,12,13).

This study aims to discover plant-derived small molecules that act as agonists and antagonists in modulating cell signaling pathways for the treatment of breast cancer and have a higher affinity to target proteins than common chemical agonists and antagonists (14,15,16). We also designed an in silico siRNA to knock down the PKMYT1 gene. In this study, a 3D-scaffold carrying these compounds is designed and introduced for local breast cancer therapy (17,18).

This study focuses on essential target proteins that play a vital role in the development and progression of breast cancer. In the following, these receptors and protein factors are introduced.

The PKMYT1 gene participates in cell cycle regulation by encoding CDK1 inhibitory kinase. In estrogen receptor-positive (ER+) breast cancer, high levels of PKMYT1 correlate with resistance to CDK4/6 inhibitors. Therefore, targeting PKMYT1 with siRNAs increases the effectiveness of inhibitors, thus potentially promoting and improving the results of resistant breast cancer treatment (19,20,21).

Aromatas: This enzyme is involved in the biosynthesis of estrogen, which causes the growth of breast cancer cells. As a result, inhibition of this enzyme reduces estrogen level, thus reducing the development of cancer (22,23,24).

Estrogen Receptor Alpha (ERα): Following the activation of this nuclear receptor by estrogen, cancer cells multiply. So, blocking this nuclear receptor inhibits tumor growth (25,26,27).

Human Epidermal Growth Factor Receptor 2 (HER2): This receptor shows a high increase in expression in a group of aggressive breast cancers. Targeted inhibition of this receptor has improved the level of treatment in HER2-positive patients (28,29,30).

Melatonin receptor 2 (MT2): MT2 plays a role in regulating cancer cell proliferation and apoptosis. Agonists of this receptor can be effective in the treatment of breast cancer (31,32,33).

Poly (ADP-ribose) polymerase 10 (PARP10): This polymerase enzyme participates in DNA repair in cancer cells. Its inhibition can increase the effectiveness of drugs that damage the DNA, and as a result, improve the efficiency of treatment (34,35,36).

Stimulator of Interferon Genes (STING): Activating the STING can activate anti-tumor innate immunity (37,38,39).

The use of natural agonists and antagonists with higher affinity than common chemical types for these receptors and enzymes leads to more targeted and successful treatments in breast cancer. In clinical studies, targeting these receptors in cancer therapy has been promising.

## 2. Methods

### 2.1. Software and Tools

The following software and online tools were used in this research. Molegro Virtual Docker (version 7): It was used to perform virtual screening and evaluate the binding affinity of selected ligands with the target receptor (40).

Molegro Molecular Viewer (version 7): It was used to visualize and analyze the results of molecular docking.

Discovery Studio 2024 Client: It was used to simulate protein-ligand interactions and to calculate RMSD (https://discover.3ds.com/discovery-studio-visualizer-download).

SwissADME: This online tool predicted ADME (absorption, distribution, and excretion) characteristics (http://www.swissadme.ch).

PDB (Protein Data Bank): Database for downloading 3D protein structures in molecular docking studies (https://www.rcsb.org).

ProTox-II: This online tool was used to analyze the toxicity of compounds (https://tox-new.charite.de/protox_II/).

siDirect 2.1: is a web-based tool, aimed at efficient and specific design of small interfering RNA (siRNA) sequences for mammalian RNA interference (RNAi). siDirect 2.1 incorporates advanced algorithms to minimize off-target effects by considering the thermodynamic stability of the duplex siRNA seed sequence (http://sidirect2.rnai.jp/).

RNAfold: This web-based tool predicts the secondary structures of single-stranded DNA or RNA sequences. It also provides both minimum free energy (MFE) structures and partition function calculations that include base pairing probabilities and ensemble diversity measures (http://rna.tbi.univie.ac.at/cgi-bin/RNAWebSuite/RNAfold.cgi).

RNAhybrid: This site-based tool predicts and calculates the binding affinity of siRNA to the target mRNA sequence. This tool performs hybridization thermodynamic calculations to determine the best binding site, and the highest certainty of gene silencing ( https://bibiserv.cebitec.uni-bielefeld.de/rnahybrid).

### 2.2. Design and evaluation of siRNA efficiency

siDirect 2.1 online tool was used to design the siRNA sequence to knock down the PKMYT1 gene. PKMYT1 gene nucleotide sequence was downloaded from NCBI RefSeq database in FASTA format, and imported into siDirect 2.1 tool. The combined laws of Ui-Tei, Reynolds and Amarzguioui were selected to design siRNA with high efficiency and high target specificity. To ensure optimal stability and performance, the GC content of siRNA was adjusted between 35% and 65%. Tm was set to 21.5 °C for maximum seed-duplex stability and minimal off-target effects. Finally, a set of potential sequences were designed to target the PKMYT1 gene, which included a guide strand with 21nt, and a complementary messenger strand with 21nt. Also, their off-target effects and thermodynamic properties were predicted.

To predict the secondary structure of PKMYT1 mRNA, the complete mRNA sequence was uploaded to the RNAfold web server. The server simulated the secondary structure using its parameters. Next, we analyzed the resulting secondary structure to identify accessible regions, such as loops and unpaired regions, required for efficient siRNA binding.

To calculate the minimum free energy (MFE) of siRNA-PKMYT1 interaction, PKMYT1 coding sequence (CDS) in FASTA format and siRNA sequence were entered into the RNAHybrid tool. After analysis based on default parameters, MFE values were calculated to evaluate the stability and efficiency of siRNA binding to target mRNA.

### 2.3. Protein Preparation

By referring to the PDB site, the 3D structures of the following proteins were prepared with a specific ID, and in PDB format: Aromatase (PDB ID: 5JL6), Estrogen Receptor Alpha (ERα) (PDB ID: 5GS4), HER2 (PDB ID: 7PCD), MT2 (PDB ID: 6ME6), PARP10 (PDB ID: 5LX6), and STING (PDB ID: 4QXQ). These structures, derived from Homo sapiens using X-ray diffraction.They have resolutions of 3.00 Å, 2.4 Å, 1.77 Å, 2.8 Å, 1.25 Å, and 2.42 Å respectively.

### 2.4. Ligand Selection

With a comprehensive literature review, a library of natural plant ligands was prepared (around 400 different compounds) using studies that determined the chemical compounds of different medicinal plants by experimental methods. The scientific names of the plants are reported in Table 4.

Then, by referring to the PubChem online database, the compounds in the prepared ligand library were searched. The 3D structure of each ligand, with its PubChem CID, was downloaded in SDF format for molecular docking study.

### 2.5. Molecular docking studies

Molegro Virtual Docker (MVD) version 7.0.0 software was used for virtual screening to find ligands with the highest binding affinity to the target proteins. The co-crystallized ligand with the crystallographic structure of the protein was used to more accurately determine the position of the active site or the binding site of the ligand in the protein structure. The position of the ligand binding sites in the structure of the proteins was also located in the places of the cavities predicted by the MVD software.

For each target protein, the grid box parameters included the following information:

Aromatase (X: 85.44, Y: 51.51, Z: 43.64, Radius: 15, Grid resolution: 0.30), ERα (X: 34.03, Y: -2.49, Z: 20.81, Radius: 11, Grid resolution: 0.30), HER2 (X: 8.76, Y: -6.97, Z: -19.15, Radius: 15, Grid resolution: 0.30), MT2 (X: 35.28, Y: -19.21, Z: 134.87, Radius: 13, Grid resolution: 0.30), PARP10 (X: 42.25, Y: 47.11, Z: 82.77, Radius: 18, Grid resolution: 0.30), and STING (X: 164.95, Y: 52.70, Z: 2.36, Radius: 25, Grid resolution: 0.30).

Molegro Virtual Docker employs several docking algorithms, including: MolDock; MolDock is a high-accuracy docking algorithm, that mimics natural selection processes in determining the optimal docking poses for ligands using a differential evolution optimization technique; MolDock SE, which is a simplified, less accurate version of MolDock, which performs high-throughput docking simulations at a faster rate; and PLP (Piecewise Linear Potential), A scoring function in combination with MolDock, which evaluates the binding affinity of ligands to proteins. In this research, the MolDock SE algorithm was used to implement the molecular docking process.

The protein structure file in PDB format was imported to MVD software. Ligands, cofactors, and water molecules were removed from the protein structure. The software identified charges and bonds and added hydrogens to the protein structure. Next, the software identified the cavities that are the potential binding sites of the ligands to the protein. Also, if necessary, the protonation states of the specific residues were set.

In the following, the SDF file of ligands downloaded from PubChem was imported. Bonds and structures of ligands were optimized by MVD. Then, energy minimization was performed to ensure stability. Then the grid box coordinates for each protein were determined as previously mentioned. Finally, a virtual screening was performed by selecting the Docking Wizard. In the end, the best pose for each ligand was determined and saved by the software.

For each protein, standard chemical agonists or antagonists were included as controls, along with the ligand library in the virtual screening process. The natural ligands that showed a higher binding affinity to the protein than the chemical agonists or antagonists were selected and evaluated for the following stages of the research.

At the stage of choosing the appropriate protein structure for molecular docking, using Discovery Studio 2024 Client, Root Mean Square Deviation (RMSD) analysis was performed to confirm the accuracy of the docking results.

### 2.6. Evaluation of CYP inhibition profiles and physicochemical properties of natural small molecules of the ligand library, and ligands of the control group

SwissADME and ProTox-III were used to evaluate the mutagenicity, cytotoxicity, and inhibitory effect of ligands on various cellular cytochrome (CYP) enzymes, including CYP1A2, CYP2C19, CYP2C9, CYP2D6, CYP3A4, and CYP2E1. Also, these online sites were used to determine physicochemical properties, molecular weight, lipophilicity (using iLOGP, XLOGP3, WLOGP, MLOGP, SILICOS-IT, and consensus), drug similarity according to Lipinski’s law, Log S (ESOL)/Class, prediction of gastrointestinal absorption (GI) and LD50. All parameters were calculated for the new small molecules in this study and the control group.

### 2.7. Proposed 3D Scaffold Design for Localized Breast Cancer Therapy

In this project, with the aim of local breast cancer therapy, a 3D scaffold is proposed to increase the delivery and effectiveness of natural small molecules. The scaffold material will be a polymer (lactic-co-glycolic acid) (PLGA), which has the characteristic of biocompatibility and gradual degradation in the physiological environment (41). This scaffold provides adequate mechanical support, and the gradual degradation is synchronized with the rate of tissue regeneration. The porous structure and large interconnected pores of the scaffold mimic well the extracellular matrix (ECM) of the breast tissue (42,43). Therefore, the scaffold structure provides a suitable microenvironment for cell growth and tissue repair by enabling cell penetration and nutrient release (44).

To maintain the stability and biological activity of natural small molecules, they are enclosed in liposomal carriers. Then, by integrating phospholipid liposomes, containing small molecules with three-dimensional scaffolds, the possibility of gradual, stable, and effective drug release in the breast area is provided (45). Scaffolds with pore sizes from 100 to 300 micrometers (µm), are suitable for breast tissue reconstruction and maintain the balance between mechanical strength and biological function.

To increase the strength of adhesion and proliferation of cells on the surface and inside the pores of the scaffold, we modify the scaffold with collagen and fibronectin (46,47). The proposed scaffold design, by maintaining the structure of small molecules, ensures their targeted and sustained release, and provides a promising approach for the in situ treatment of breast cancer.

One of the advantages of this scaffold design is its ability in local treatment, reducing systemic side effects and improving the therapeutic effectiveness of small molecules. Controlled release and bioavailability of small molecules is possible by integrating liposomes carrying them in the scaffold structure. In designing this innovative approach, the advantages of 3D tissue engineering and targeted drug delivery have been used. As a result, it provides an effective solution for breast cancer treatment.

## 3. Resultes

### 3.1. Insilico analysis of siRNA

A particular siRNA was designed to target the PKMYT1 gene, and targeted the CDS nucleotide sequences at positions 400-422 (Or, 5’-836, 816-3’, on the whole RNA sequence) of the PKMYT1 gene. The guide strand with the sequence ACGCUUUACCGCAUAGAGCCG, which was complementary to the target mRNA strand, established a precise binding with the target strand (21nt target + 2nt overhang: CGGCTCTATGCGGTAAAGCGTTC/3’-816,838-5’). This precise binding leads to the destruction of the mRNA. The passenger strand with the sequence, GCUCUAUGCGGUAAAGCGUUC (3’-818,838-5’) was complementary to the guide strand. This complementarity, results forming a stable duplex before being processed by the RNA-induced silencing complex (RISC).

In efficient formation of the RISC complex, seed-duplex stability is essential, the guide strand and the passenger strand provided such conditions with melting temperatures (Tm) of 20.4 °C and 17.8 °C, respectively. Analysis showed minimal off-target effects; there, were two potential off-target ( Homo sapiens holocarboxylase synthetase (HLCS), and Homo sapiens agouti signaling protein (ASIP)) hits with one mismatch and three hits with two mismatches for the guide strand, whereas for the passenger strand, no off-target hits with 0 or There was one mismatch, but two hits with three mismatches. Therefore, the designed siRNA is particular for the PKMYT1 gene and reduces the possibility of off-target effects.

As a result, the designed siRNA with minimal off-target effects and the ability to form a stable duplex was recognized as a suitable candidate for PKMYT1 knockdown.

#### 3.1.1. Position of the Target Sequence

In siRNA design, accessibility to the mRNA target sequence is essential for effective siRNA binding. In the complex structure of mRNA, the target sequence of siRNA can be out of reach. By predicting the mRNA secondary structure, it is possible to identify the accessible regions of the mRNA located inside the loop structures and not inside the stem. Paying attention to this point increases the efficiency of the siRNA and reduces off-target effects. In the secondary structure predicted for PKMYT1 mRNA, the position of the target sequence was nucleotides 816 to 836, and part of the sequence was in the loop structure.

The result of the Rnahybrid analyzer showed that the siRNA guide sequence targets the PKMYT1 mRNA (NM_182687.3) with a minimum free energy (MFE) of -45.1 kcal/mol, which indicates a strong binding affinity. The alignment of siRNA and target sequence had a stable interaction, Fig.1.

**Fig. 1.**
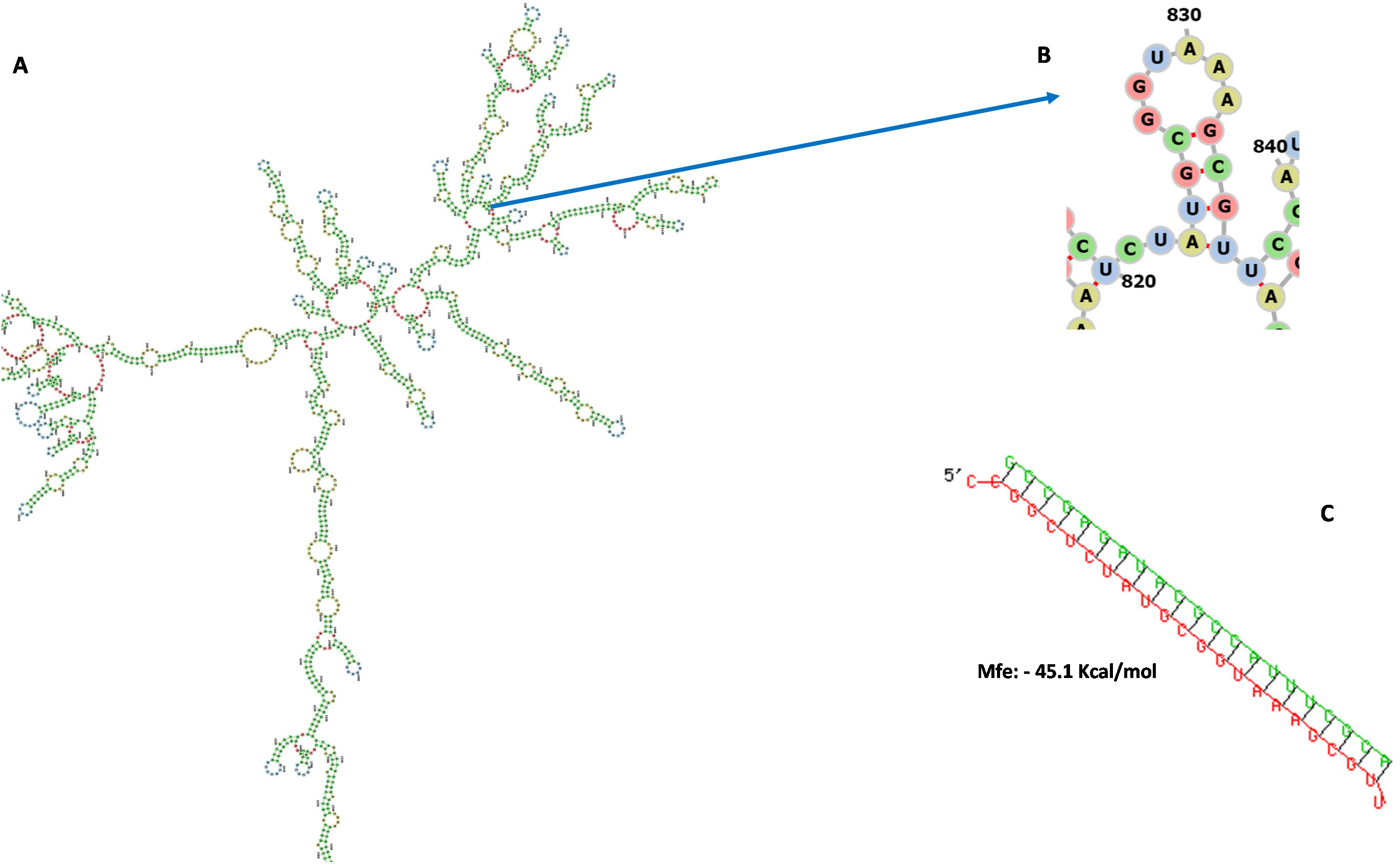
Two-dimensional structure predicted for PKMYT1 mRNA by RNAfold web server(A). siRNA target site in PKMYT1 Mrna (B). Calculation of binding affinity of siRNA to the target site in mRNA by RNAhybrid (C).

**Fig. 2.**
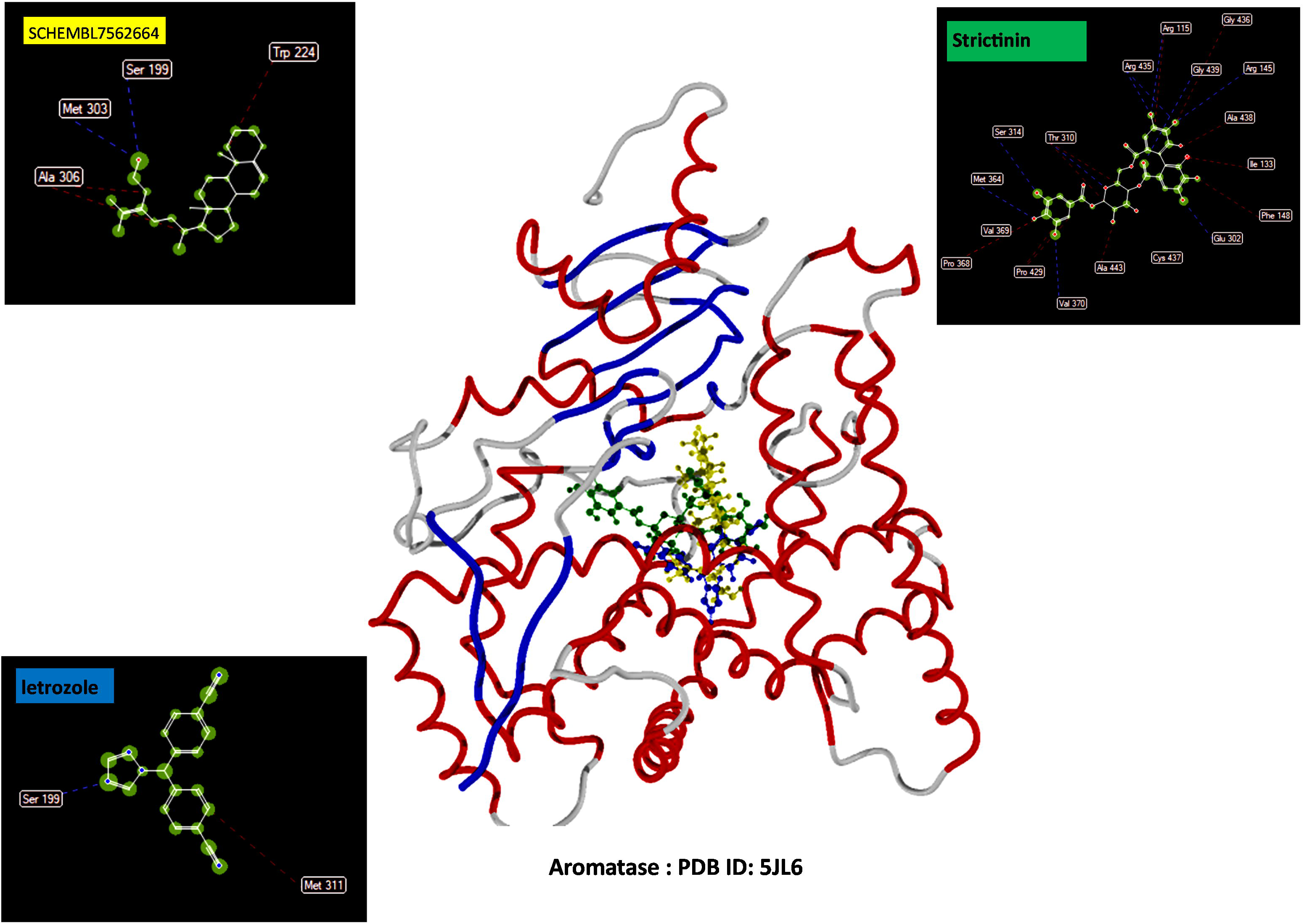
Visualization of ligand-Aromatase, chemical interactions: control ligand (blue label), golden ligand (yellow label) and ligand with the highest binding affinity (green label).

**Fig. 3.**
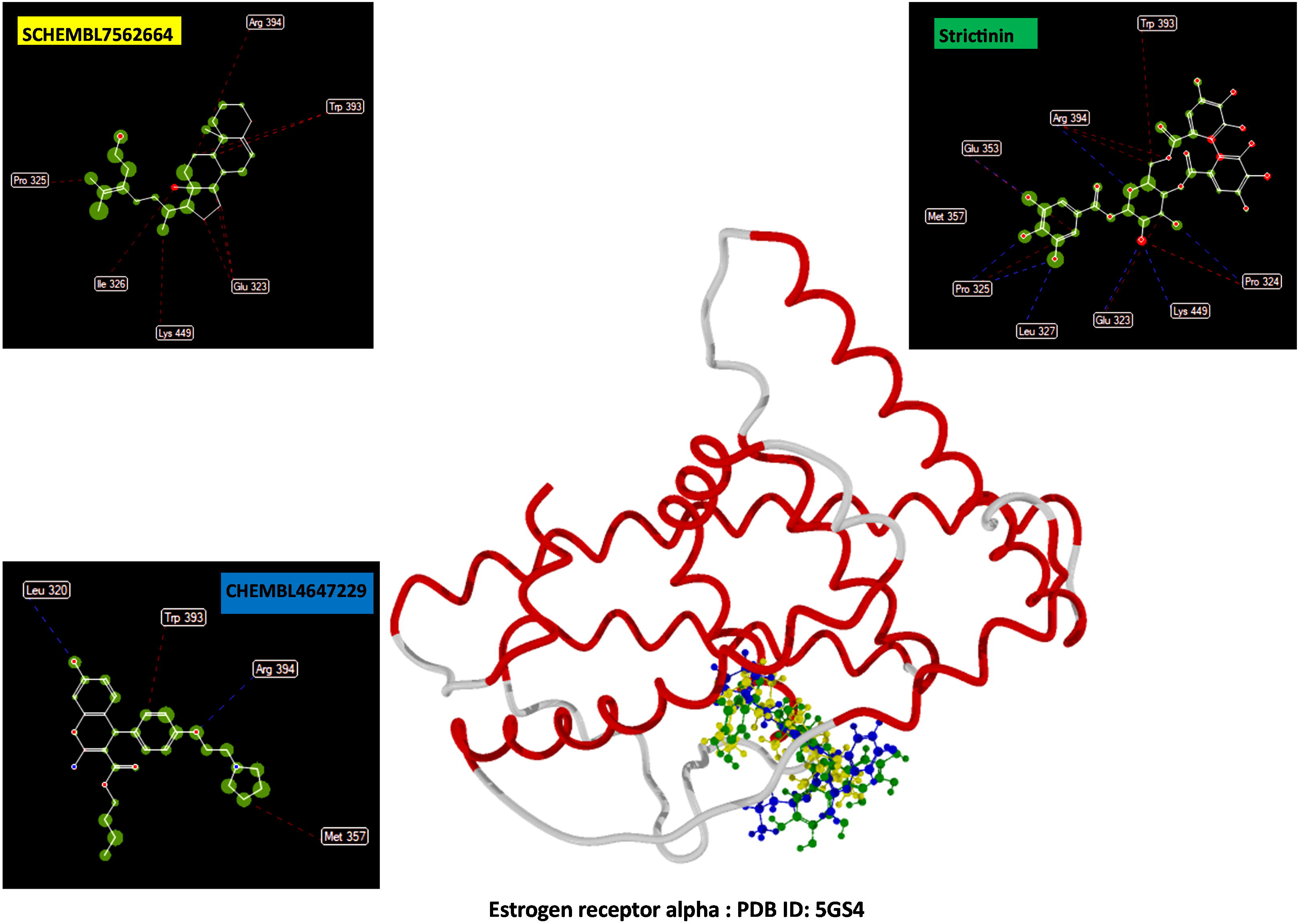
Visualization of ligand-Estrogen receptor alpha, chemical interactions: control ligand (blue label), golden ligand (yellow label) and ligand with the highest binding affinity (green label).

**Fig. 4.**
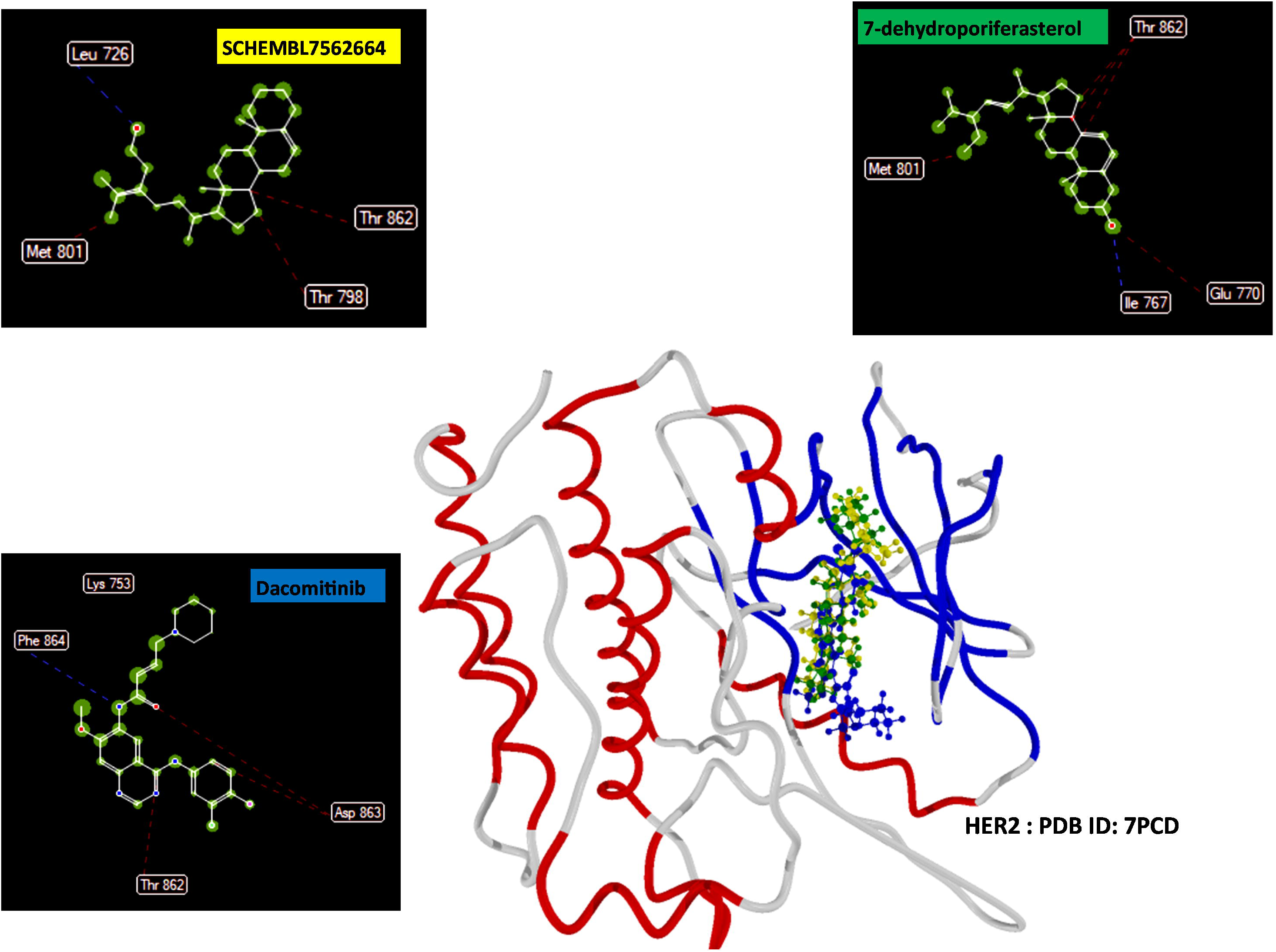
Visualization of ligand-HER2, chemical interactions: control ligand (blue label), golden ligand (yellow label) and ligand with the highest binding affinity (green label).

**Fig. 5.**
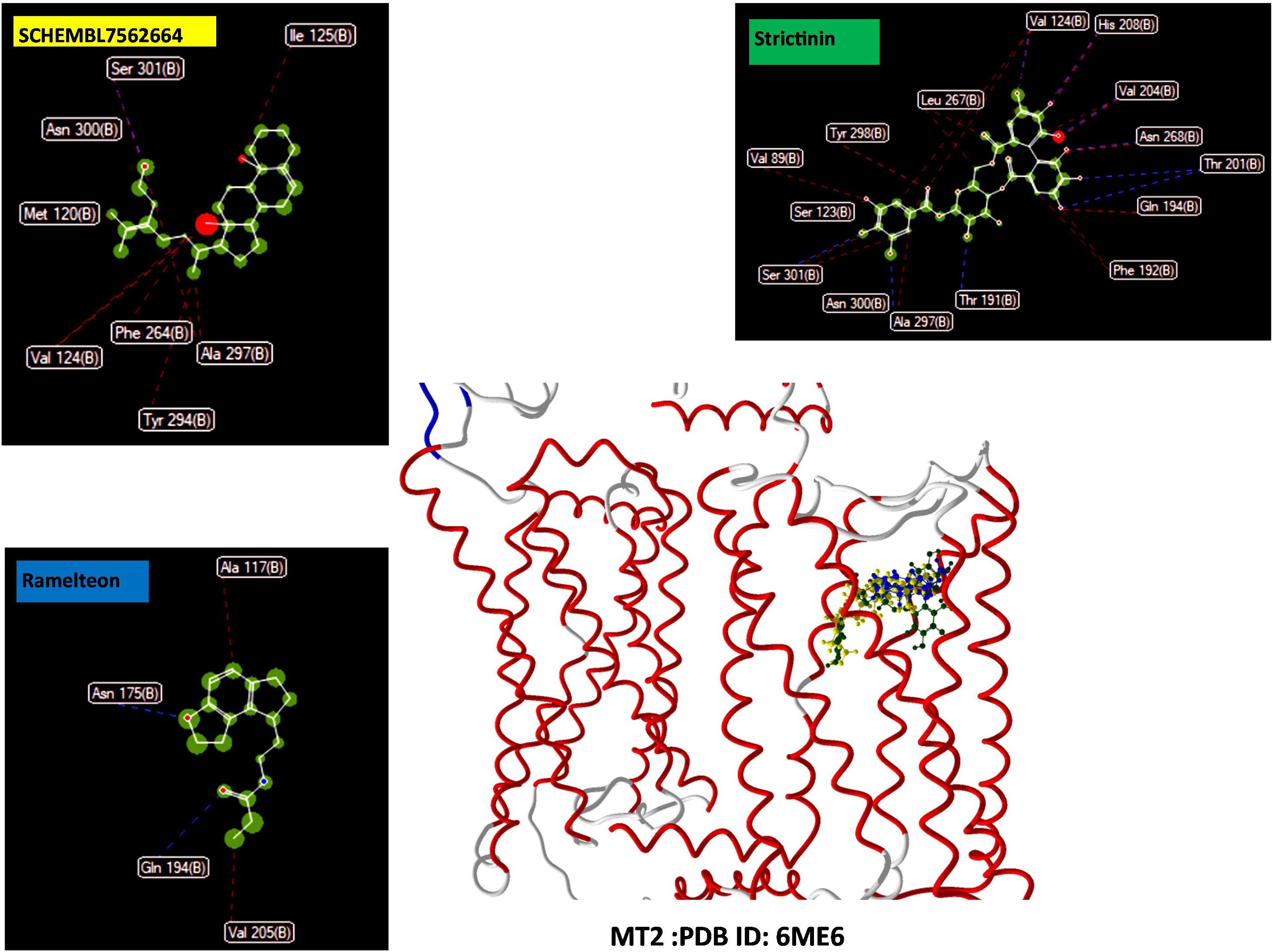
Visualization of ligand-MT2, chemical interactions: control ligand (blue label), golden ligand (yellow label) and ligand with the highest binding affinity (green label).

**Fig. 6.**
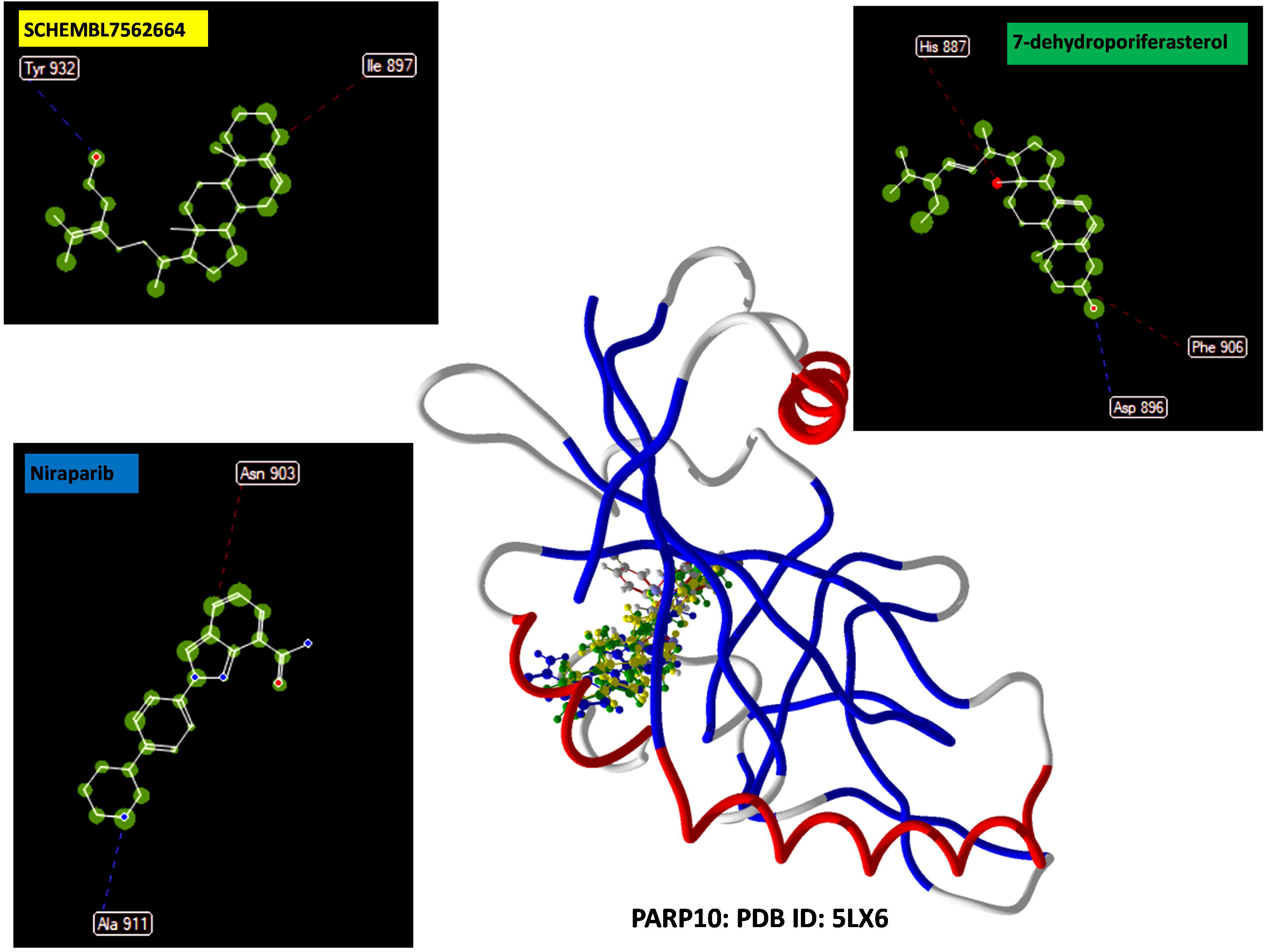
Visualization of ligand-PARP10, chemical interactions: control ligand (blue label), golden ligand (yellow label) and ligand with the highest binding affinity (green label).

**Fig. 7.**
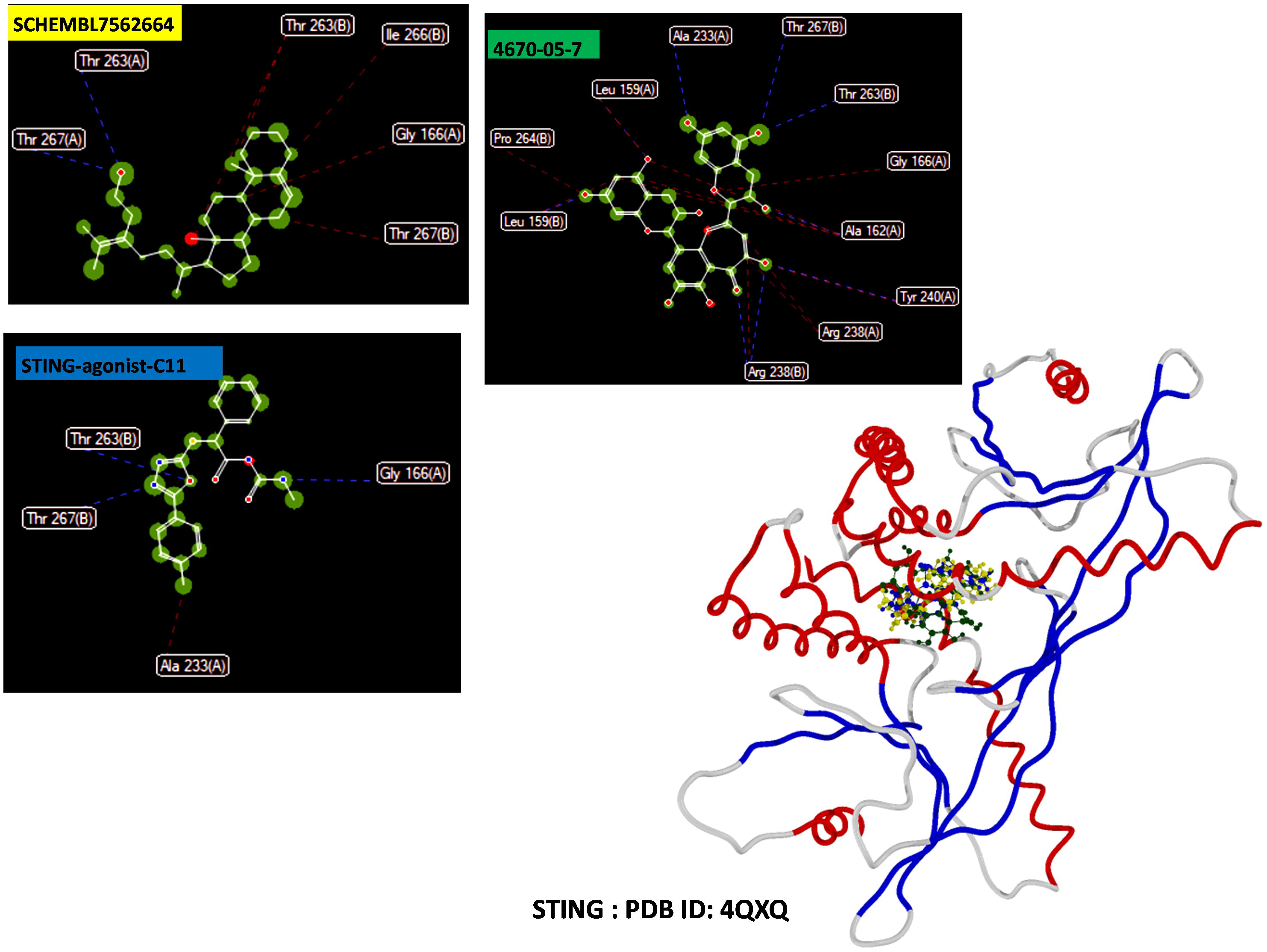
Visualization of ligand-STING chemical interactions: control ligand (blue label), golden ligand (yellow label) and ligand with the highest binding affinity (green label).

### 3.2. Analysis of binding affinity of selected ligands with their target proteins

Binding affinity, expressed by the equilibrium dissociation constant (Kd), indicates and evaluates the strength of the interaction between a ligand (such as a drug or a molecule) and its target protein. More negative Kd values correspond to higher binding affinity, meaning the ligand binds more tightly to its target protein (49).

1. The binding affinity of Strictinin to aromatase (PDB ID: 5JL6) was -161.941 kJ/mol, which was significantly higher than the control ligands; Letrozole (-110.042 kJ/mol), Aromasin (-105.63 kJ/mol), and Testolactone (-97.4206 kJ/mol). Its affinity to estrogen receptor alpha (PDB ID: 5GS4), was -129.202 kJ/mol, higher than Tamoxifen (-111.095 kJ/mol), and CHEMBL4647229 (-117.612 kJ/mol). For HER2 (PDB ID: 7PCD), it showed -116.78 kJ/mol binding affinity, which was lower than Lapatinib (-143.425 kJ/mol), and Neratinib (-125.739 kJ/mol), but higher than Dacomitinib (-114.611 kJ/mol). For MT2 (PDB ID: 6ME6), its binding affinity was -170.097 kJ/mol, which was, much higher than Ramelteon (-112.932 kJ/mol). For PARP10 (PDB ID: 5LX6), it was -135.075 kJ/mol, lower than Talazoparib (-153.637 kJ/mol) and Olaparib (-146.085 kJ/mol), but higher than Niraparib (-107.072 kJ/mol). Strictinin did not have binding affinity to the STING (PDB ID: 4QXQ).

2. Glansreginin A showed a binding affinity of -158.16 kJ/mol to the aromatase, higher than Letrozole, Aromasin, and Testolactone. This ligand did not have binding affinity to estrogen receptor alpha, HER2, and PARP10. For MT2, its binding affinity (-150.348 kJ/mol), was higher than Ramelteon. Also, it was - 118.735 kJ/mol, for STING, lower than STING-agonist-C11 (-146.381 kJ/mol), and STING agonist-1 (- 126.657 kJ/mol), but higher than STING agonist-14 (-106.02 kJ/mol).

3. 5’-Methoxyhydnocarpin had a binding affinity of -142.396 kJ/mol for aromatase, higher than Letrozole, Aromasin, and Testolactone, but did not bind to estrogen receptor alpha. For HER2, it showed -124.091 kJ/mol, lower than Lapatinib and Neratinib, but higher than Dacomitinib. For MT2, it was - 126.55 kJ/mol, higher than Ramelteon. It was -108.86 kJ/mol, for PARP10, which was lower than Talazoparib, but higher than Niraparib. It did not show binding affinity to the STING.

4. SCHEMBL9475140 exhibited a binding affinity of -132.533 kJ/mol for aromatase, higher than Letrozole, Aromasin, and Testolactone. But did not show binding affinity to estrogen receptor alpha, and HER2. For MT2, this ligand showed -128.993 kJ/mol binding affinity, higher than Ramelteon. For PARP10, it was -147.532 kJ/mol, higher than Olaparb, and Niraparib, but lower than Talazoparib. For STING, was -133.747 kJ/mol, lower than STING-agonist-C11 but higher than STING agonist-1, and STING agonist-14.

5. D5-Avenasterol acetate had a binding affinity of -130.868 kJ/mol for aromatase, higher than Letrozole, Aromasin, and Testolactone. But did not show binding affinity to estrogen receptor alpha, and HER2. For MT2, It showed a binding affinity of -141.348 kJ/mol, higher than Ramelteon. For PARP10, it was -140.355 kJ/mol, higher than, Niraparib, lower than Talazoparib and Olaparb. For STING, it showed - 137.361 kJ/mol, lower than STING-agonist-C11 but higher than STING agonist-1, and STING agonist-14.

6. Epicatechin-(4beta-6)-epicatechin showed a binding affinity of -128.661 kJ/mol for aromatase, higher than Letrozole, Aromasin, and Testolactone, but did not have binding affinity to estrogen receptor alpha, HER2 and MT2. For PARP10, its binding affinity was -148.931 kJ/mol, higher than Olaparb and Niraparib, lower than Talazoparib. It was -119.042 kJ/mol for STING, lower than STING-agonist-C11, and STING agonist-1, but higher than STING agonist-14.

7. GAMA-TOCOPHEROL exhibited a binding affinity of -124.466 kJ/mol for aromatase, higher than Letrozole, Aromasin, and Testolactone, but did not show binding affinity to estrogen receptor alpha. For HER2, it had a binding affinty of -116.765 kJ/mol, lower than Lapatinib,and Neratinib, but higher than Dacomitinib. For MT2, it was -143.189 kJ/mol, higher than Ramelteon. It showed a binding affiniy of, - 130.135 kJ/mol, For PARP10, it was higher than Niraparib but lower than Talazoparib, and Olaparib. It had a binding affinity of -137.386 for STING, lower than STING-agonist-C11, higher than STING agonist-1, and STING agonist-14.

8. Elaidic acid showed a binding affinity of -107.248 kJ/mol for aromatase, lower than Letrozole, but highter than Aromasin, and Testolactone. For estrogen receptor alpha, it showed -111.659 kJ/mol, almost equal to Tamoxifen, and lower than CHEMBL4647229. For MT2, it had a binding affinity of - 121.443 kJ/mol, higher than Ramelteon. For PARP10, it was -121.793 kJ/mol, lower than Talazoparib and Olaparib, higher than Niraparib. And for STING, it showed a binding affinity of -118.202 kJ/mol, lower than STING-agonist-C11, and higher than STING agonist-14. It did not show binding affinity to HER2.

9. 7-dehydroporiferasterol had a binding affinity of -115.811 kJ/mol for aromatase, higher than Letrozole, Aromasin, and Testolactone. For estrogen receptor alpha, it did not show binding affinity. For HER2 it had a binding affinity of -129.879 kJ/mol, higher than Neratinib, and Dacomitinib. For MT2, it showed -150.931 kJ/mol, higher than Ramelteon. For PARP10, it was -149.976 kJ/mol, higher than Olaparib, and Niraparib, lower than Talazoparib. For STING, it showed a binding affinity of -153.122 kJ/mol, higher than STING control ligands.

10. 4670-05-7 demonstrated a binding affinity of -153.102 kJ/mol for aromatase, higher than Letrozole, Aromasin, and Testolactone. It did not have a binding affinity for estrogen receptor alpha. For HER2, it showed -120.971 kJ/mol, higher than Dacomitinib, but lower than Lapatinib, and Neratinib. For MT2, it showed a binding affinity of -136.177 kJ/mol, higher than Ramelteon. For PARP10, it was -133.324 kJ/mol, higher than Neratinib, but lower than Olaparib and Talazoparib. For STING, it had a binding affinity of -160.91 kJ/mol, higher than STING control ligands.

11. SCHEMBL7562664 had a binding affinity of -114.939 kJ/mol for aromatase, higher than Letrozole, Aromasin, and Testolactone. For estrogen receptor alpha, it showed a binding affinity of -115.746 kJ/mol, higher than Tamoxifen and lower than CHEMBL4647229. For HER2, it was -119.279 kJ/mol, lower than Lapatinib and Neratinib, but higher than Dacomitinib. For MT2, it was -141.849 kJ/mol, higher than Ramelteon, and for PARP10, it showed a binding affinity of -128.661 kJ/mol, lower than Olaparib and Talazoparib, but higher than Niraparib. For STING, it was -147.469 kJ/mol, higher than STING control ligands.

12. Clerosterol exhibited a binding affinity of -113.138 kJ/mol for aromatase, higher than Letrozole, Aromasin, and Testolactone. It did not have a binding affinity For estrogen receptor alpha. It showed a binding affinity of -117.664 kJ/mol for HER2, higher than Dacomitinib, but lower than Lapatinib, and Neratinib. For MT2, it was -131.592 kJ/mol, higher than Ramelteon. For PARP10, it was -123.951 kJ/mol, higher than Niraparib, lower than Olaparib and Talazoparib. For STING, it was -152.067 kJ/mol, higher than STING control ligands.

13. 24-Ethylcholesterol had a binding affinity of -117.654 kJ/mol for aromatase, higher than Letrozole, Aromasin, and Testolactone. It did not show binding affinity for estrogen receptor alpha, and HER2. For MT2, it was -139.357 kJ/mol, higher than Ramelteon. For PARP10, it was -131.561 kJ/mol, higher than Niraparib, but lower than Olaparib and Talazoparib. For STING, it showed a binding affinity of -151.859 kJ/mol, higher than STING control ligands.

All results related to binding affinity of selected and control ligands to target proteins are reported in Table 1.

**Table 1.** In this table, the binding affinity of different natural ligands to crucial proteins involved in breast cancer treatment is summarized. Each ligand is self-identified by number, name, PubChem CID, and molecular formula. The table includes data on the ligand’s affinity (in kJ/mol) for aromatase (PDB ID: 5JL6), estrogen receptor alpha (PDB ID: 5GS4), HER2 (PDB ID: 7PCD), MT2 (PDB ID: 6ME6), PARP10 (PDB ID: 5LX6), and STING (PDB ID: 4QXQ). The top three ligands with the highest binding affinity for each target protein are highlighted. SCHEMBL7562664 has been introduced as a golden ligand due to its exceptional multi-targeting capabilities and lack of cytotoxic and mutagenic effects. Control ligands are also included for comparison. Ligands with lower binding affinity than all corresponding control ligands to the specific target protein are considered to have no significant binding affinity for the target protein.

### 3.3. CYP Inhibition Profiles, Mutagenicity, and Cytotoxicity

The inactivation of cellular cytochromes (CYPs) by small molecules and drugs plays a crucial role in maintaining metabolic balance and detoxification processes in the body. Among the results of inhibiting these enzymes by small molecules and drugs are adverse drug interactions, accumulation of toxins, and general disruption in metabolic processes. In addition, it is essential that the ligands mustn’t have any cytotoxic or mutagenic effects, thus preventing small molecules from causing cell damage, cancer, and other serious health problems (50,51). The results of the evaluation for all selected ligands and control are listed in Table 2.

**Table 2.** This table summarizes the inhibitory effects of different ligands on different cytochrome P450 enzymes (CYP1A2, CYP2C19, CYP2C9, CYP2D6, CYP3A4, and CYP2E1), as well as their mutagenicity and cytotoxicity. Each specific ligand is evaluated for its activity, and compared with control ligands.

5’-Methoxyhydnocarpin was active against CYP2C19, CYP2C9, and CYP3A4. SCHEMBL9475140 was active against CYP2C19 and CYP2C9. 7-dehydroporiferasterol was active against CYP2C9. All three ligands of SCHEMBL7562664, Clerosterol, and 24-Ethylcholesterol were active against CYP2C9. All these ligands were without mutagenic or cytotoxic effects.

The other ligands, including Strictinin, Glansreginin A, D5-Avenasterol acetate, Epicatechin-(4beta-6)-epicatechin, GAMA-TOCOPHEROL, Elaidic acid, and 4670-05-7, were inactive against all CYPs and did not show cytotoxic or mutagenic effects.

Among the ligands in the control group, Letrozole was active against CYP1A2 and CYP3A4. Aromasin was active for CYP2C9. Tamoxifen was active for CYP2C9 and CYP3A4. Lapatinib was active against CYP2C9, CYP2D6, and CYP3A4, and also was cytotoxic. Dacomitinib was active against CYP2C9 and CYP2D6, this ligand was mutagenic. Ramelteon was active against CYP2D6. Olaparib was active for CYP2C9. Niraparib was active against CYP2D6. STING agonist-1 was active against CYP2C9. STING agonist-14 was active for CYP1A2, CYP2C19, CYP2C9, CYP2D6, and CYP3A4, also, also was mutagenic.

Other control ligands were inactive against CYPs and without mutagenic and cytotoxic effects.

### 3.4. Druglikeness, Log S (ESOL)/Class, GI Absorption, and Predicted LD50

Table (3) shows the chemical properties and biological characteristics of selected ligands and control ligands. Key concepts include: hydrogen bond donors and acceptors, determining the ability of each ligand to form hydrogen bonds, which affects solubility and interaction with biological targets.

The molecular weight of ligands affects their ability to penetrate the cell membrane. Lipophilicity (Log Po/w) indicates the degree of solubility of a ligand in lipids versus water, which involves the absorption and distribution of a specific ligand in the body. Drug similarity, which is determined by Lipinski’s law, evaluates the possibility of using the ligand as an oral drug. Solubility (Log S) affects the ligand’s ability to dissolve in body fluids, and its bioavailability. GI absorption predicts the extent of ligand absorption in the gastrointestinal tract. LD50 estimates the lethal dose for 50% of the population, and represents the degree of toxicity of each ligand.

The molecular weight of Strictinin was 634.45 g/mol, and it had three violations of Lipinski’s law, which can affect its pharmaceutical similarity. Its moderate lipophilicity (Log Po/w Consensus 0.76) confirmed its balanced hydrophilicity and lipophilicity. This compound was soluble (Log S: -3.92) but had a low GI absorption that could limit its oral bioavailability. The predicted LD50 was 2260 mg/kg, indicating relatively low toxicity. Glansreginin A also had three violations of Lipinski’s law, with a molecular weight of 593.58 g/mol. It had moderate lipophilicity (Log Po/w Consensus 0.12), was also soluble (Log S: -2.38), and showed low GI absorption and LD50, 2000 mg/kg.

5’-Methoxyhydnocarpine conformed to Lipinski’s law and showed no violations. Its molecular weight was 494.45 g/mol, and had high lipophilicity (consensus Log Po/w 2.66). This ligand was moderately soluble (Log S: -5.36). It had low GI absorption and predicted LD50 of 5000 mg/kg, indicating low toxicity. SCHEMBL9475140 had two violations of Lipinski’s law and a molecular weight of 586.89 g/mol. It showed very high lipophilicity (consensus Log Po/w 9.23), so it was insoluble (Log S: -10.93), with low GI absorption and a predicted LD50 of 2510 mg/kg.

D5-Avenasterol acetate had one violation of Lipinski’s Rule with a molecular weight of 468.75 g/mol. It had high lipophilicity (consensus Log Po/w of 7.70). It was poorly soluble (Log S: -8.43), with low GI absorption and a predicted LD50 of 1185 mg/kg. Epicatechin-(4beta-6)-epicatechin had three violations of Lipinski’s Rule, a molecular weight of 578.52 g/mol, and moderate lipophilicity (consensus Log Po/w of 1.42). It was moderately soluble (Log S: -5.14), with low GI absorption and a predicted LD50 of 2500 mg/kg.

GAMA-TOCOPHEROL had one violation of Lipinski’s Rule, with a molecular weight of 416.68 g/mol, and high lipophilicity (consensus Log Po/w of 7.94). It was poorly soluble (Log S: -8.29). And showed low GI absorption, and a predicted LD50 of 5000 mg/kg. Elaidic acid showed one violation of Lipinski’s Rule, a molecular weight of 282.46 g/mol, and high lipophilicity (consensus Log Po/w of 5.71). It was moderately soluble (Log S: -5.41), with high GI absorption, also a predicted LD50 of 48 mg/kg, which indicated higher toxicity.

7-dehydroporiferasterol had one violation of Lipinski’s Rule, with molecular weight of 410.68 g/mol, and high lipophilicity (consensus Log Po/w of 6.81). It was poorly soluble (Log S: -7.08) with low GI absorption, also a predicted LD50 of 10 mg/kg, which indicated higher toxicity. 4670-05-7 showed three violations of Lipinski’s Rule, a molecular weight of 564.49 g/mol, and moderate lipophilicity (consensus Log Po/w of 1.07). It was moderately soluble (Log S: -5.12). And had low GI absorption, with a predicted LD50 of 2500 mg/kg.

SCHEMBL7562664 had one violation of Lipinski’s Rule, with a molecular weight of 412.69 g/mol, also high lipophilicity (consensus Log Po/w of 7.26). It was poorly soluble (Log S: -7.96) with low GI absorption, and a predicted LD50 of 640 mg/kg. Clerosterol showed one violation of Lipinski’s Rule, with molecular weight of 412.69 g/mol, and high lipophilicity (consensus Log Po/w of 7.19). It was poorly soluble (Log S: -7.84) and had low GI absorption, with a predicted LD50 of 890 mg/kg. 24-Ethylcholesterol had one violation of Lipinski’s Rule, a molecular weight of 414.71 g/mol, and high lipophilicity (consensus Log Po/w of 7.23). It was poorly soluble (Log S: -7.90) also, showed low GI absorption, with a predicted LD50 of 890 mg/kg.

The control ligands, such as Letrozole, Aromasin, and Testolactone, Generally were consistent with Lipinski’s law. They also showed moderate to high lipophilicity, good solubility, high GI absorption, and different toxicity levels. For example, Letrozole, had a molecular weight of 285.3 g/mol, no violations of Lipinski’s Rule, moderate lipophilicity (consensus Log Po/w of 2.32), and was soluble (Log S: -3.70) with high GI absorption and a predicted LD50 of 1463 mg/kg. Aromasin and Testolactone showed similar profiles.

### 3.5. Protein-Ligand Binding Affinity Analysis

The top three ligands with the highest binding affinity for each receptor are introduced below.

For Aromatase (PDB ID: 5JL6), the ligands with the highest affinity were Strictinin (-161.941 kJ/mol), Glansreginin A (-158.16 kJ/mol), and 4670-05-7 (-153.102 kJ/mol). For Estrogen Receptor Alpha (PDB ID: 5GS4), the top ligands were Strictinin (-129.202 kJ/mol), SCHEMBL7562664 (- 115.746 kJ/mol), and Elaidic acid (-111.656 kJ/mol), In the case of HER2 (PDB ID: 7PCD), the highest affinities were shown by 7-dehydroporiferasterol (-129.879kJ/mol), 5’-Methoxyhydnocarpin (-124.091 kJ/mol), and 4670-05-7 (- 120.971 kJ/mol).

For MT2 (PDB ID: 6ME6), the ligands with the highest affinity were Strictinin (-170.097 kJ/mol), 7-dehydroporiferasterol (-150.931 kJ/mol), and Glansreginin A (-150.348 kJ/mol). For PARP10 (PDB ID: 5LX6), the top ligands were 7-dehydroporiferasterol (-149.976 kJ/mol), Epicatechin-(4beta-6)-epicatechin (-148.931 kJ/mol), and SCHEMBL9475140 (-147.532 kJ/mol). For STING (PDB ID: 4QXQ), the ligands with the highest affinity were 4670-05-7 (-160.91 kJ/mol), 7-dehydroporiferasterol (-153.122 kJ/mol), and Clerosterol (-152.067 kJ/mol).

### 3.6. Golden ligand in this study, and new ligands for target proteins investigated in this study

Among the selected ligands in this in silico study, SCHEMBL7562664 could to target all six, target proteins simultaneously. Based on the data in tables 1, 2 and 3, It’s affinity for each receptor was higher than the affinity of some control ligands for each target protein. This compound did not have any cytotoxic and mutagenic effects. And it was only active against CYP2C9. Due to the high lipophilicity it showed, it was designated as a golden candidate for breast cancer local therapy. Also, based on the literature review, this ligand is introduced for the first time for all the proteins examined in this research Fig. 8-12.

**Table 3.** This table provides a detailed overview of various ligands, focusing on their hydrogen bond donors and acceptors, molecular weight, lipophilicity, drug-likeness according to Lipinski’s Rule, solubility, gastrointestinal absorption, and predicted lethal dose (LD50). Each ligand’s properties are listed to help assess their potential as drug candidates.

**Table 4.** This table lists the scientific names of plants whose constituent compounds were experimentally determined. These compounds were used to create a ligand library for virtual screening.

**Fig. 8.**
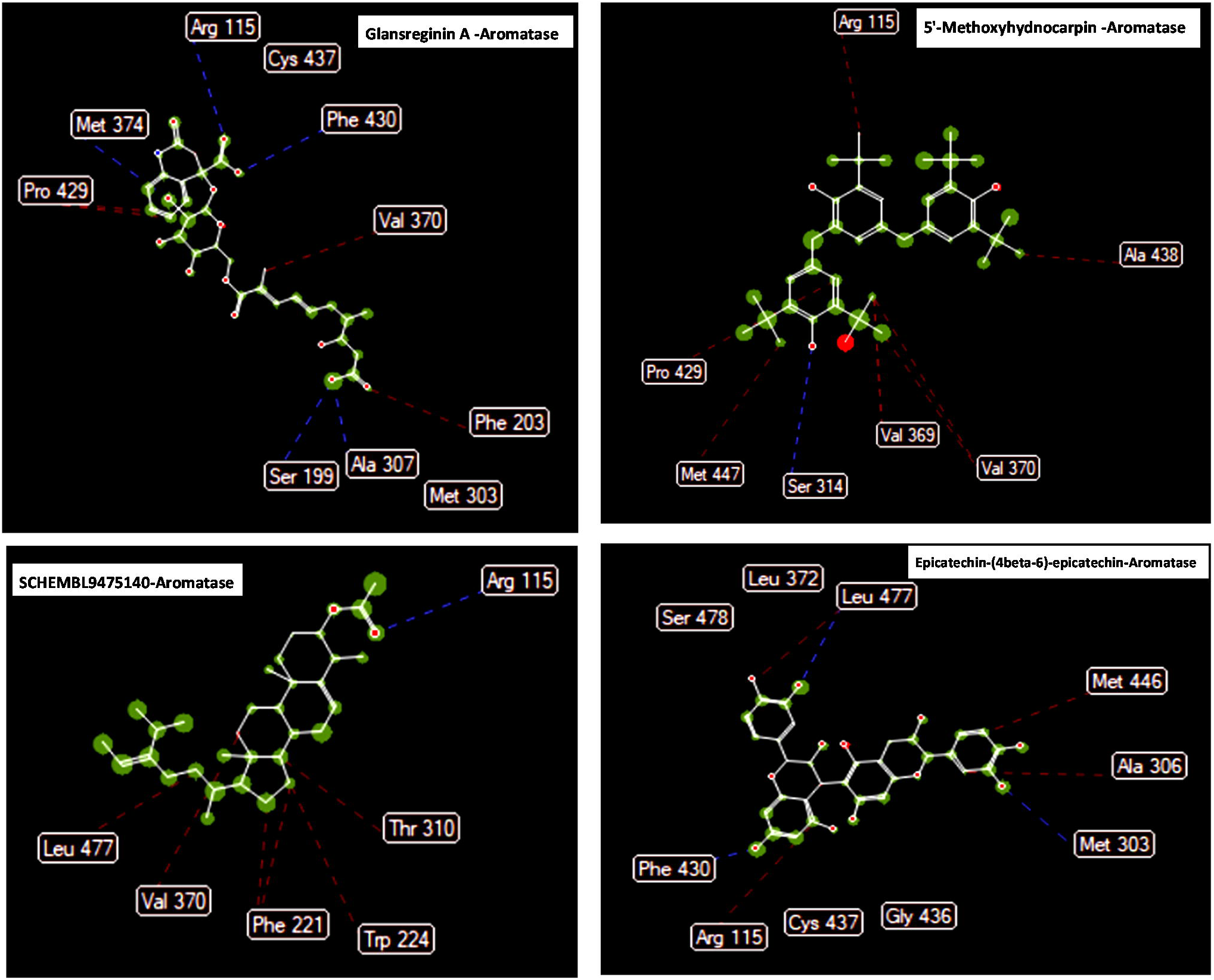
Ligand map related to new antagonists based on the results of the present study and literature review, with Aromatase. Hydrogen bonds are shown with blue dashed lines and steric interactions with red dashed lines.

**Fig. 9.**
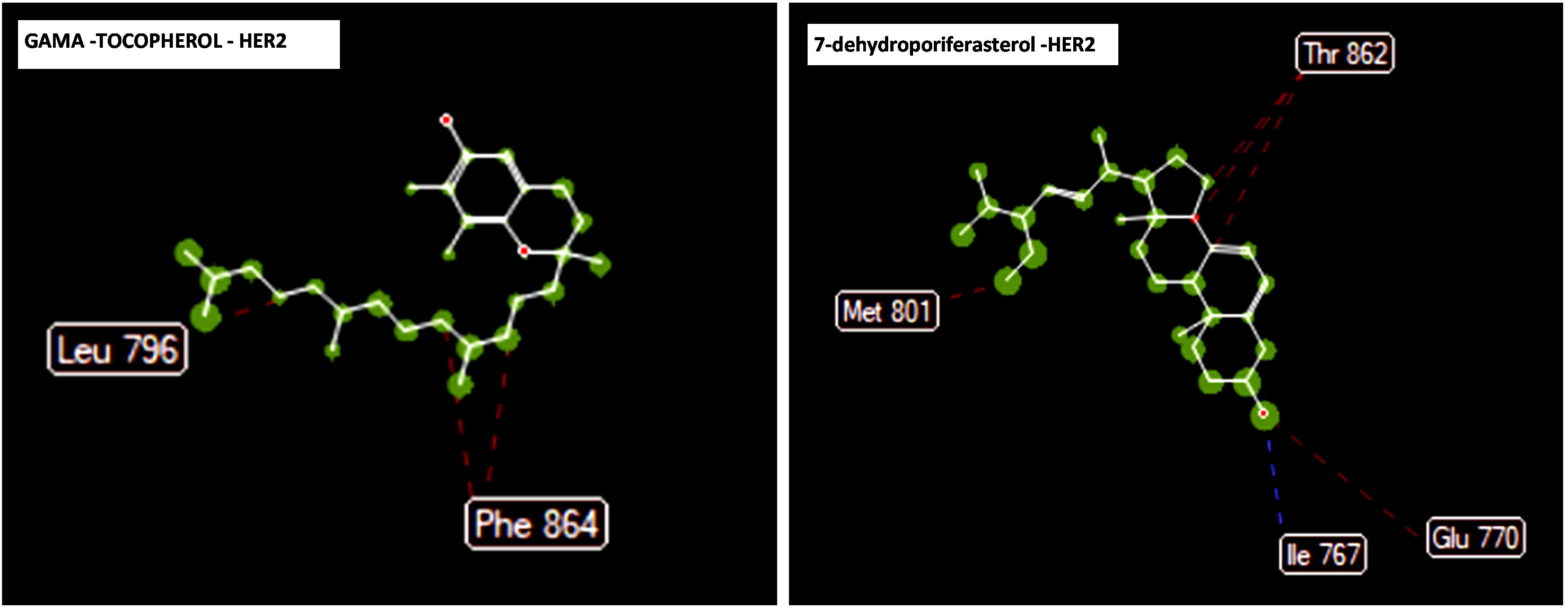
Ligand map related to new antagonists based on the results of the present study and literature review, with HER2. Hydrogen bonds are shown with blue dashed lines and steric interactions with red dashed lines.

**Fig. 10.**
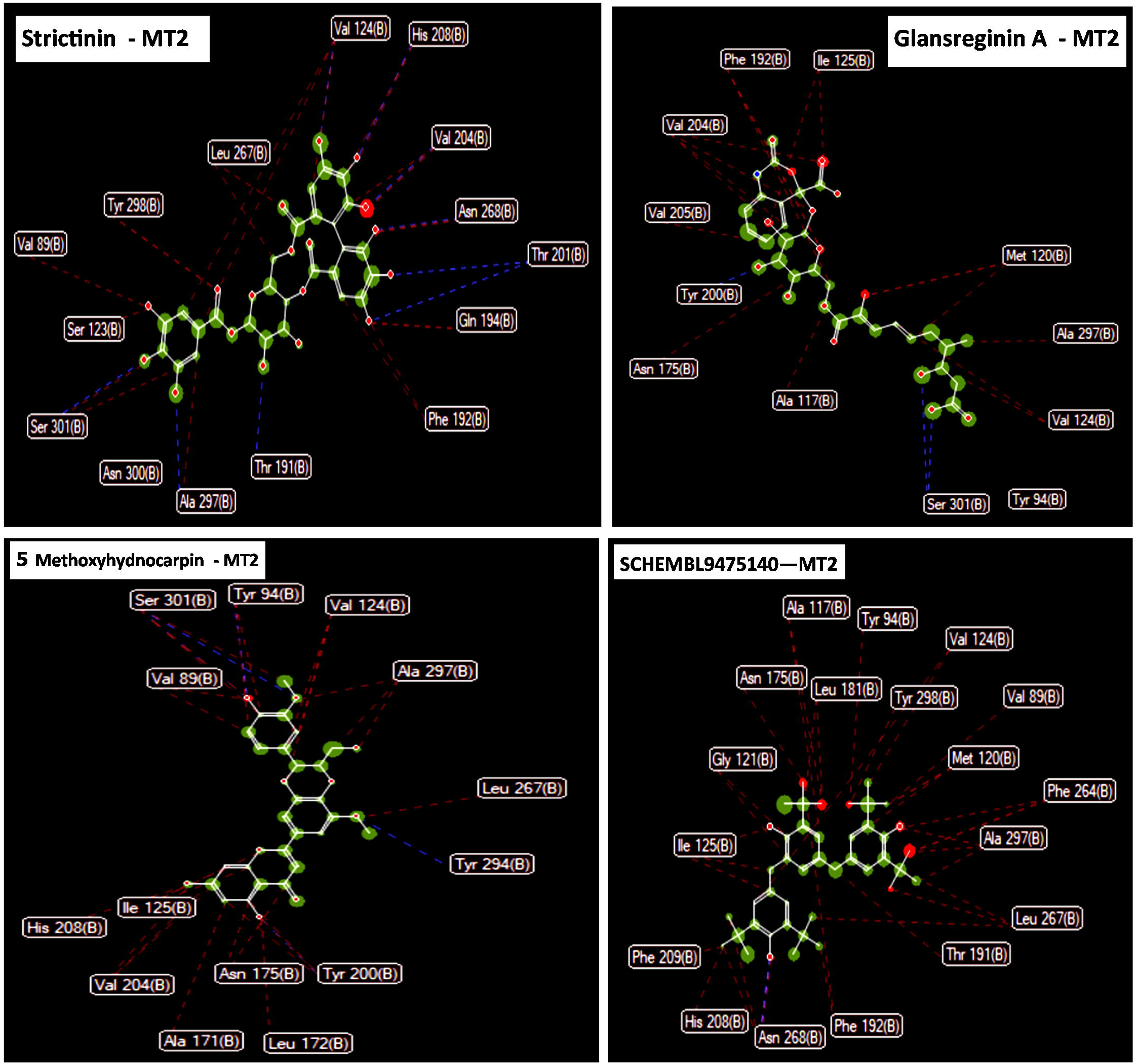
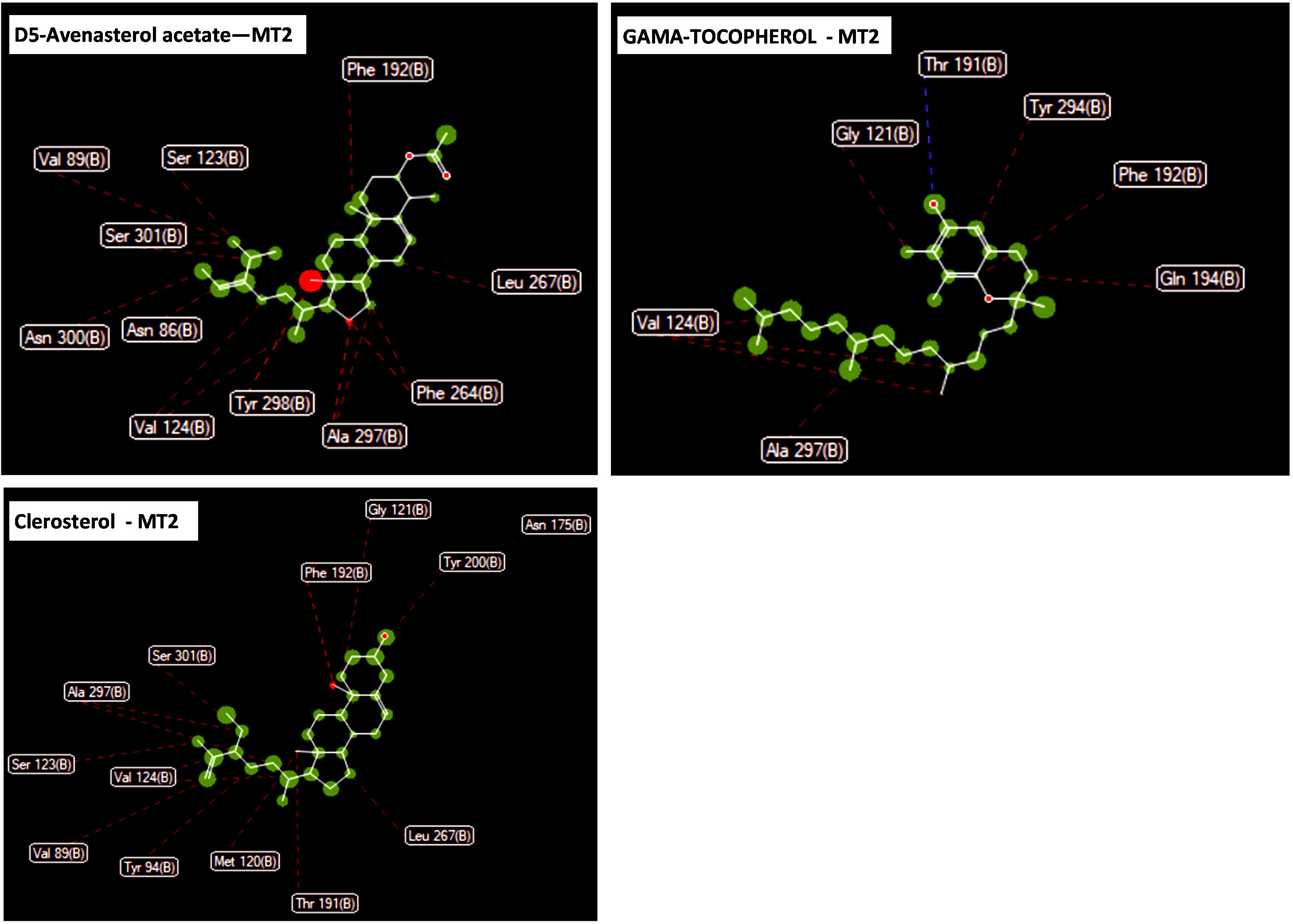
**(a and b)** Ligand map related to new agonists based on the results of the present study and literature review, with MT2 protein. Hydrogen bonds are shown with blue dashed lines and steric interactions with red dashed lines.

**Fig. 11.**
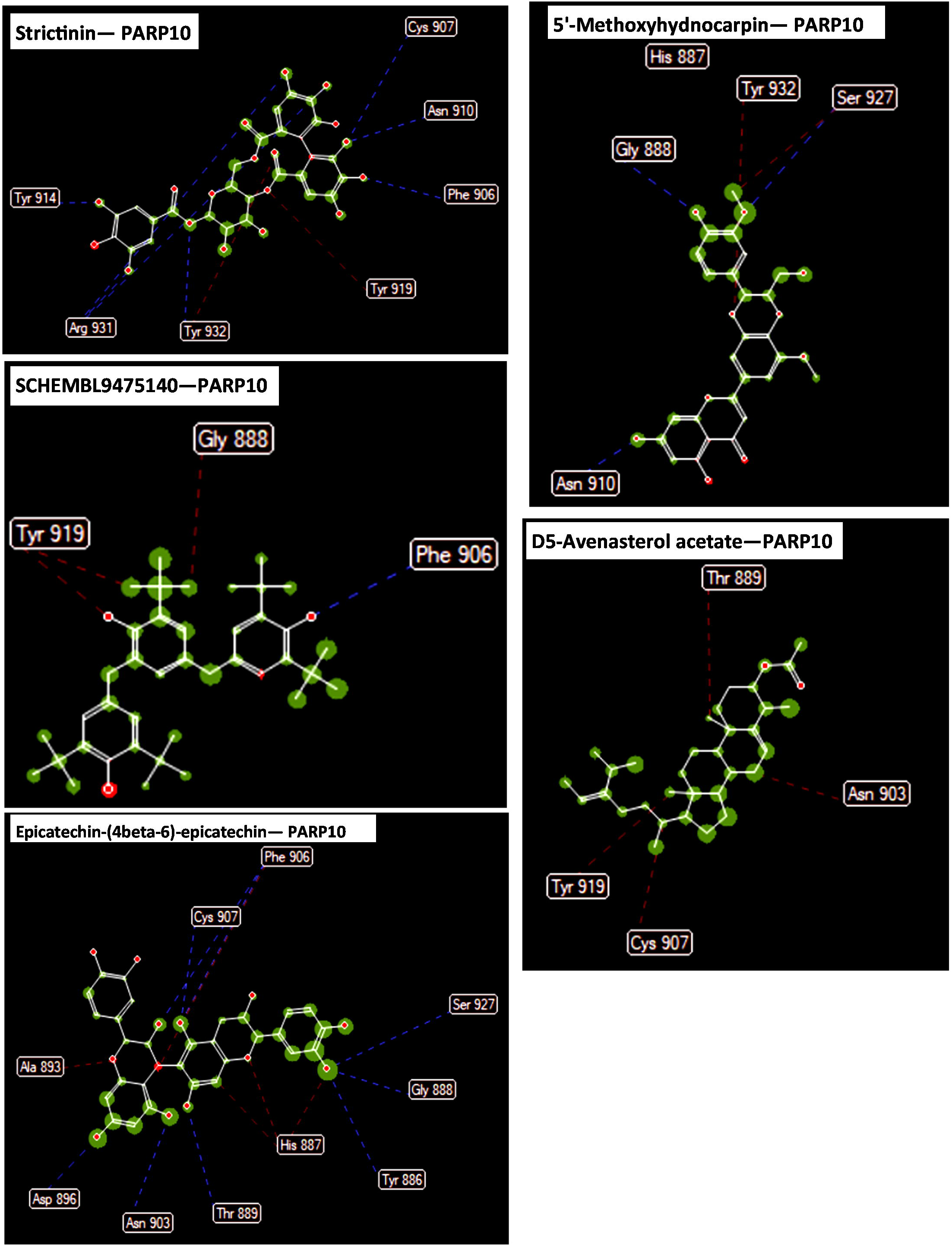
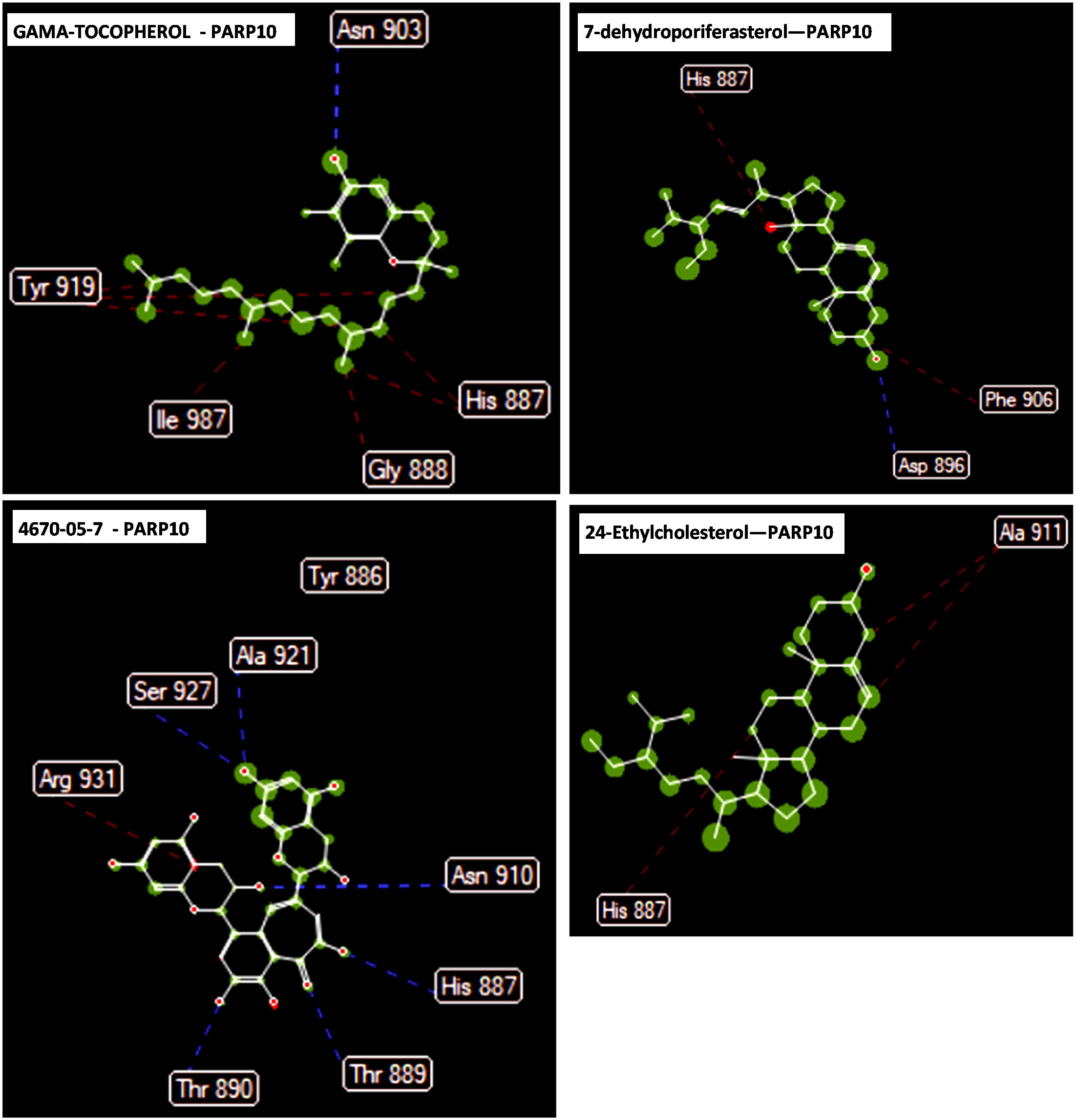
**(a and b)** Ligand map related to new antagonists based on the results of the present study and literature review, with PARP10. Hydrogen bonds are shown with blue dashed lines and steric interactions with red dashed lines.

**Fig. 12.**
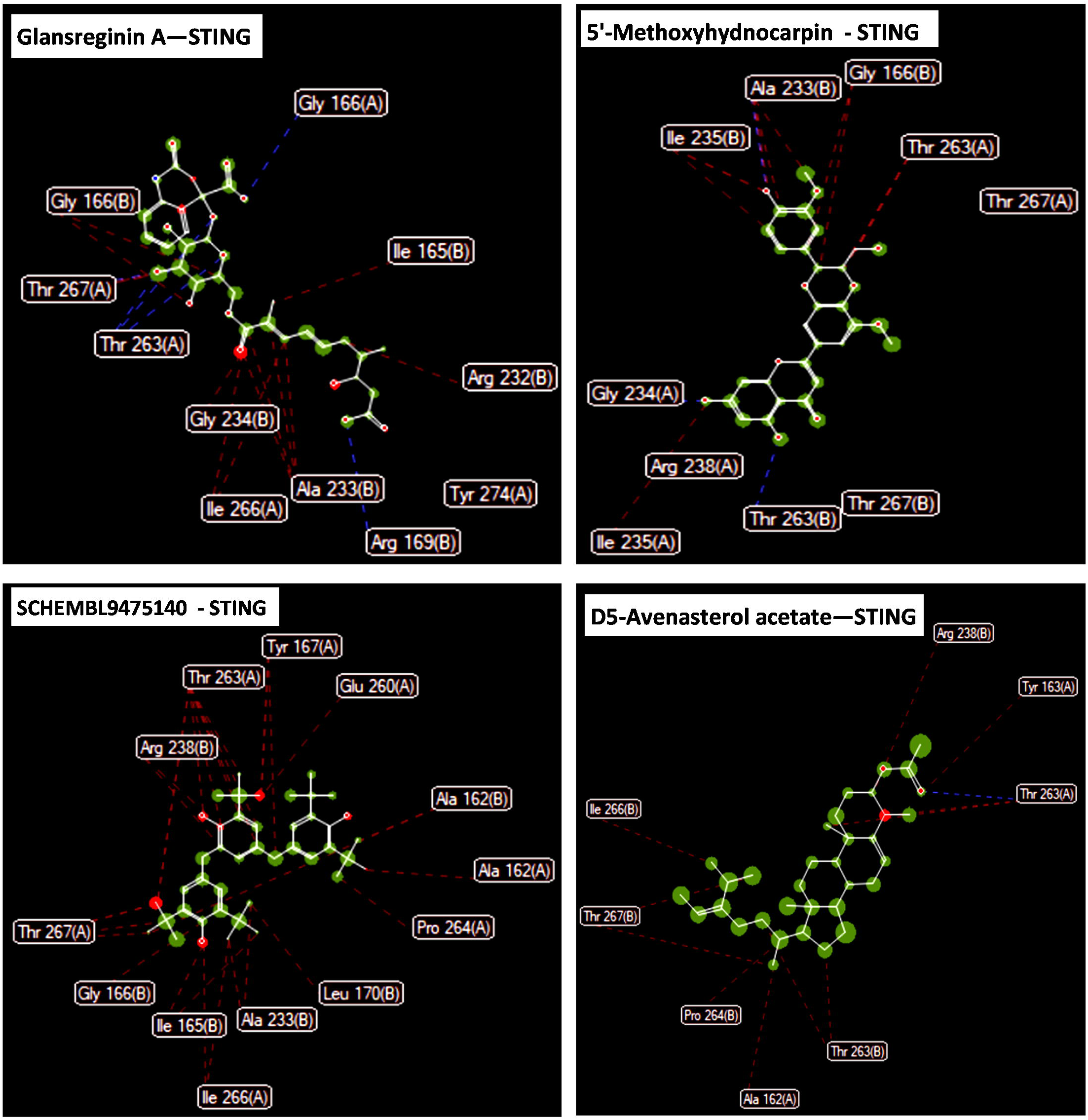
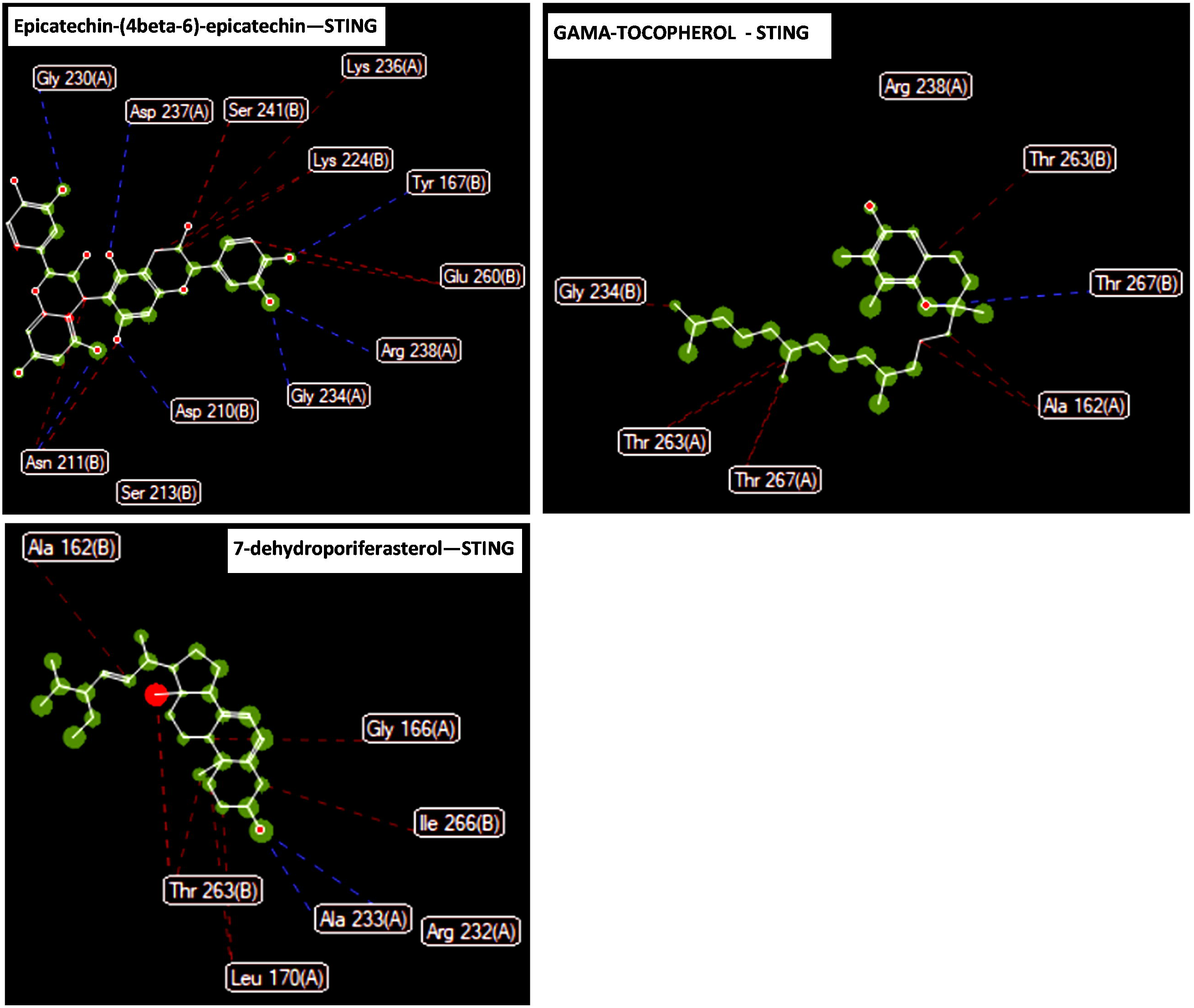
**(a and b)** Ligand map related to new agonists based on the results of the present study and literature review, with STING protein. Hydrogen bonds are shown with blue dashed lines and steric interactions with red dashed lines.

The source of all selected ligands in this study is listed in Table 6.

**Table 5.** In this table, the origin of the selected ligands from the plants used in this study is determined.

**Table 6.** In this table, the calculated RMSD values for each protein are listed.

### 3.7. Root Mean Square Deviation (RMSD) analysis

Root Mean Square Deviation (RMSD) is a measure used to quantify the difference between values predicted by a model and the values observed. It is also used in structural biology to compare the three-dimensional structures of proteins and nucleic acids. RMSD values less than 2 angstroms (Å), are considered acceptable, indicating a high degree of similarity between the compared structures. RMSD is used to measure the accuracy of the predicted binding positions of ligands to their receptors in molecular docking. If the RMSD value is less than 2 Å, it suggests that the predicted pose closely matches the experimentally determined pose , Table 7.

Based on data in Table 4, the RMSD values for Aromatase, ERa, HER2, MT2, PARP10, and STING were below 2 Å.

### 3.8. Visualization of ligand-protein interactions

Using Molegro Molecular Viewer software, version 7, the interactions between the ligand and the amino acids in the binding site in the protein structure were accurately drawn. The protein backbone is represented by the tube model, while the ligands are color-coded for clarity: the colors are blue for the control ligand, yellow for the golden ligand, and green for the ligand with the highest binding affinity. The ligand map shows the chemical interaction of each ligand with a specific residue in the binding site, hydrogen bonds are shown as dashed blue lines and steric interactions are shown as dashed red lines. The ligand maps are for three golden ligand, the ligand with higher affinity, and the control ligand, the name of each ligand is written with yellow, blue, and green color labels in the corresponding ligand map. Protein chain names are shown in capital letters, which provide a detailed and comprehensive representation of binding interactions, Fig 2-7.

### 4. Discussion

In this study, in order to increase the delivery quality and effectiveness of natural small molecules in the local therapy of breast cancer, the design of a 3D scaffold made of poly(lactic-co-glycolic acid) (PLGA) was proposed. PLGA, having biocompatibility and gradual degradation in physiological environments, is compatible with the speed of tissue regeneration. The scaffold, with its porous structure composed of interconnected pores, from 100 to 300 µm, closely ECM of breast tissue, thus providing mechanical support and a favorable microenvironment for cell growth and tissue repair. The combination of liposomal carriers inside the scaffold causes the stability and biological activity of small molecules and their controlled and stable release in the breast tissue. The purpose of using this scaffold is to minimize systemic side effects and improve therapeutic effectiveness by maintaining a high concentration of therapeutic agents in the tumor site, in the breast tissue.

In previous researches in the local therapy of breast cancer using 3D scaffolds, promising results in increasing drug delivery and efficiency have been reported. For example, the study of Abuwatfa et al., highlighted the benefits of using scaffolds in cancer research, also demonstrating the ability of 3D scaffolds to mimic the tumor microenvironment, and improve drug sensitivity (52).

Our study builds on these findings from other researchers’ studies, along with the incorporation of liposomal carriers into the PLGA scaffold, which not only mimics the ECM, but also ensures a controlled and sustained release of therapeutic agents.

siRNA designed to target the PKMYT1 gene, with high specificity and minimal off-target effects, became a promising candidate for gene knockdown. The guide strand makes precise binding with a complementary sequence of the target mRNA, causing mRNA degradation. The passenger strand forms a stable duplex with the guide strand, which is necessary for efficient formation of the RISC complex. The predicted secondary structure of PKMYT1 mRNA showed that the target sequence is accessible to siRNA. The strong binding affinity, with a minimum free energy (MFE) of -45.1 kcal/mol, confirmed the potential of this siRNA for effective gene silencing.

Previous research highlights the potential of PKMYT1 knockdown as a therapeutic target in breast cancer. For example, Chen et al., confirmed that PKMYT1 is a therapeutic response marker and a target for CDK4/6 inhibitor-resistant ER+ breast cancer (53). They showed that the use of specific inhibitors such as lunresertib (RP-6306), combined with gemcitabine, selectively knocked down PKMYT1, which resulted in reduced survival of resistant ER+ breast cancer cells.

Similarly, Lee et al., discovered that overexpression of PKMYT1 was associated with poor prognosis and immune infiltration in triple-negative breast cancer (TNBC) (54). They showed that knockdown of PKMYT1 inhibits proliferation, migration and invasion of TNBC cells, and induces apoptosis, these observations confirmed that targeting PKMYT1 can improve treatment outcomes for TNBC patients.

Our in silico analysis of siRNA targeting PKMYT1 is consistent with these findings. Designed siRNA can reduce tumor growth and improve treatment outcomes in breast cancer.

Equilibrium dissociation constant (Kd) is used to evaluate the strength of interaction between ligand and target protein. More negative Kd values correspond to higher binding affinity, and indicate tighter binding of the ligand to the target protein.

Aromatase is an enzyme that is involved in the conversion of androgens into estrogens, stimulating the growth of breast cancer cells with positive hormone receptors. In postmenopausal women, to reduce the growth of these cancer cells, aromatase inhibitors are used to reduce estrogen levels. Previous studies have identified several potent aromatase inhibitors, such as letrozole and exemestane, which are widely used in the treatment of breast cancer (55,56,57).

However, our study introduces new inhibitors such as Strictinin (-161.941 kJ/mol) and Glansreginin A (- 158.16 kJ/mol), which show much higher binding affinities compared to these established inhibitors. As a result, the inhibitors introduced in this study could potentially provide more effective inhibition of aromatase, thus leading to better therapeutic outcomes.

More than 70% of breast cancers express the nuclear transcription factor ERα. This transcription factor plays a role in regulating the transcription of genes involved in cell proliferation and tumor progression, so it has become a vital target in endocrine therapies (58). Tamoxifen is recognized as the main drug in the treatment of estrogen receptor-positive breast cancer (59). Our research highlights Strictinin (- 129.202 kJ/mol) and SCHEMBL7562664 (-115.746 kJ/mol) as novel antagonists with higher binding affinities than tamoxifen (-111.095 kJ/mol). These new antagonists can lead to improved efficacy in the treatment of breast cancer by providing stronger inhibition of ERα.

HER2 is a protein effective in cell growth. HER2 overexpression is present in approximately 15-20% of breast cancers, which is associated with aggressive tumor growth and poor prognosis (60). HER2-targeted therapies, using inhibitors such as Trastuzumab and Lapatinib, have significantly improved treatment in HER2-positive breast cancer patients (61). Our study identifies 7-dehydroporiferasterol (- 129.879 kJ/mol), 5’-Methoxyhydnocarpin (-142.396 kJ/mol), and SCHEMBL7562664 (-119.279 kJ/mol) as novel antagonists with similar or higher binding affinities than some existing HER2 inhibitors. Therefore, these new ligands can increase the therapeutic options in HER2-positive breast cancer.

MT2 plays an important role in protecting against oxidative stress. Its expression is related to tumor growth, differentiation and drug resistance (62). The potential of melatonin receptor agonists in the treatment of breast cancer has been investigated due to their role in regulating the circadian rhythm and cell proliferation. In the present study, Strictinin (-170.097 kJ/mol) and Glansreginin A (-150.348 kJ/mol) were introduced as new agonists with higher binding affinity than existing agonists like Ramelteon (63).

PARP10 has a role in DNA repair and plays an important role in maintaining genomic stability. PARP10 inhibition is associated with increased DNA damage and cancer cell death, particularly in breast cancer with BRCA mutations (64, 65). One of the available PARP inhibitors is olaparib, which has a promising effect in the treatment of breast cancer. In our research, ligands such as 7-dehydroporiferasterol (- 149.976 kJ/mol) and SCHEMBL9475140 (-147.532 kJ/mol) were introduced as new antagonists with higher binding affinities than some current PARP inhibitors, such as Olaparib and Niraparib (66).

Therefore, these novel ligands can induce enhanced inhibition of PARP10, improving treatment outcomes for patients with BRCA-mutated cancers.

As an important protein in innate immunity, STING plays an important role in antitumor immunity. Activation of the STING pathway increases the immune system’s ability to target and destroy cancer cells. STING agonists have been investigated for their role in enhancing antitumor immunity (67). Our study introduces compounds such as 7-4670-05-7 (-160.91 kJ/mol) and 7-dehydroporiferasterol (- 153.122 kJ/mol) as new agonists with higher binding affinities than existing agonists for STING. These new ligands can enhance the immune response against the tumor, and provide new strategies for cancer immunotherapy.

Among the selected ligands, SCHEMBL7562664 was introduced as an outstanding candidate because it can target all six target proteins simultaneously. This ligand showed higher binding affinity for each receptor compared to some of their control ligands, and also lacked any cytotoxic and mutagenic effects. These features make it a promising candidate for local treatment of breast cancer. Based on literature review, SCHEMBL7562664 is introduced for the first time for all the proteins investigated in this research.

This study addressed the challenge of drug resistance in breast cancer by introducing new and multi-targeted natural ligands, as well as specific siRNA sequences to inhibit multiple cancer pathways. The proposed 3D scaffold for local therapy promotes delivery, targeted release, and enhancement of the effectiveness of these agents at the tumor site. The potential outcome of this approach is to improve treatment outcomes and minimize systemic side effects. It also provides a promising strategy to overcome drug resistance in breast cancer.

Due to the in silico nature of this research, complete in vivo and in vitro validation is necessary to confirm the presented findings. Experimental investigations are needed to investigate the potential off-target effects of siRNA design and variation in ligand binding affinities under biological conditions. In addition, the activity of some ligands against cytochrome P450 enzymes can lead to drug interactions that require detailed interaction studies. To ensure the efficacy and safety of the proposed 3D scaffold for local delivery, clinical evaluations using organoid models should be performed. A careful examination of these limitations through validation and rigorous experimental testing will strengthen the study’s results and promote the development of effective breast cancer treatments.

## 5. Conclusion

In this study, new high-affinity ligands for essential target proteins involved in breast cancer were successfully identified, also, effective siRNA with minimal off-target effect for PKMYT1 gene knockdown and a three-dimensional scaffold for local delivery of these therapeutic agents were designed.

The identified ligands showed potent and selective inhibitory effects on aromatase, estrogen receptor alpha, HER2, and PARP10, as well as agonistic effects on MT2 and STING, which enhance antitumor immunity and modulate cancer progression pathways. Designed siRNA provided a precise approach to inhibit cancer cell proliferation, metastasis, and drug resistance. Also, the compound SCHEMBL7562664, was introduced as a golden ligand, because in this study it targeted all the target proteins with high affinity, and without showing cytotoxic and mutagenic effects.

The proposed 3D scaffold ensures effective delivery and controlled release of therapeutic compounds directly to the tumor site, resulting in increased treatment efficacy and minimized systemic side effects.

The multi-objective approach presented in this research not only improves the efficiency of treatment, but also reduces the possibility of developing drug resistance and provides a promising strategy for breast cancer treatment.

## Supporting information

binding affinity of selected ligands

CYP Inhibition Profiles, Mutagenicity, and Cytotoxicity ,Druglikeness

