## Supplementary material for "In Silico Discovery of Multi-Target Natural Ligands and Efficient siRNA Design for Overcoming Drug Resistance in Breast Cancer via Local Therapy": binding affinity of selected ligands

| **Ligand Number** | **Ligand Name** | **PubChem CID** | **Molecular Formula** | **Aromatase antagonist**  **PDB ID: 5JL6**  **+Ligand affinity**  **(kj/mol** | **Estrogen receptor alpha antagonist**  **PDB ID: 5GS4**  **+ Ligand affinity**  **(kj/mol** | **HER2 antagonist**  **PDB ID: 7PCD**  **+ Ligand affinity**  **(kj/mol** | **MT2 agonist**  **PDB ID: 6ME6**  **+ Ligand affinity**  **(kj/mol** | **PARP10**  **Antagonist**  **PDB ID: 5LX6**  **+ Ligand affinity**  **(kj/mol** | **STING agonist**  **PDB ID: 4QXQ**  **+ Ligand affinity**  **(kj/mol** |
| --- | --- | --- | --- | --- | --- | --- | --- | --- | --- |
| **1** | Strictinin | 73330 | C_27_H_22_O_18_ | **Yes**  **-161.941** | **Yes**  **-129.202** | Yes  -116.78 | **Yes**  **-170.097** | Yes  -135.075 | No |
| **2** | Glansreginin A | 16104376 | C_28_H_35_NO_13_ | **Yes**  **-158.16** | No | No | **Yes**  **-150.348** | No | Yes  -118.735 |
| **3** | 5'-Methoxyhydnocarpin | 5281879 | C_26_H_22_O_10_ | Yes  -142.396 | No | **Yes**  **-124.091** | Yes  -126.55 | Yes  -108.86 | Yes  -140.849 |
| **4** | SCHEMBL9475140 | 545456 | C_40_H_58_O_3_ | Yes  -132.533 | No | No | Yes  -128.993 | **Yes**  **-147.532** | Yes  -133.747 |
| **5** | D5-Avenasterol acetate | 91753899 | C_32_H_52_O_2_ | Yes  -130.868 | No | No | Yes  -141.348 | Yes  -140.355 | Yes  -137.361 |
| **6** | Epicatechin-(4beta-6)-epicatechin | 131752343 | C_30_H_26_O_12_ | Yes  -128.661 | No | No | No | **Yes**  **-148.931** | Yes  -119.042 |
| **7** | GAMA-TOCOPHEROL | 45356270 | C_28_H_48_O_2_ | Yes  -124.466 | No | Yes  -116.765 | Yes  -143.189 | Yes  -130.135 | Yes  -137.386 |
| **8** | Elaidic acid | 637517 | C_18_H_34_O_2_ | Yes  -107.248 | **Yes**  **-111.659** | No | Yes  -121.443 | Yes  -121.793 | Yes  -118.202 |
| **9** | 7-dehydroporiferasterol | 20843308 | C_29_H_46_O | YES  -115.811 | No | **Yes**  **-129.879** | **Yes**  **-150.931** | **Yes**  **-149.976** | **Yes**  **-153.122** |
| **10** | 4670-05-7 (THEAFLAVINE) | 135403798 | C_29_H_24_O_12_ | **Yes**  **-153.102** | No | **Yes**  **-120.971** | Yes  -136.177 | Yes  -133.324 | **Yes**  **-160.91** |
| **11** | SCHEMBL7562664 | 70130285 | C_29_H_48_O | Yes  -114.939 | **Yes**  **-115.746** | Yes  -119.279 | Yes  -141.849 | Yes  -128.661 | Yes  -147.469 |
| **12** | Clerosterol | 5283638 | C_29_H_48_O | Yes  -113.138 | No | Yes  -117.664 | Yes  -131.592 | Yes  -123.951 | **Yes**  **-152.067** |
| **13** | 24-Ethylcholesterol | 9823110 | C_29_H_50_O | -117.654 | No | No | Yes  -139.357 | Yes  -131.561 | Yes  -151.859 |
| **14. Aromatase antagonist control** | letrozole | 3902 | C_17_H_11_N_5_ | Yes  -110.042 |  |  |  |  |  |
| **15. Aromatase antagonist control** | Aromasin | 60198 | C_20_H_24_O_2_ | Yes  -105.63 |  |  |  |  |  |
| **16. Aromatase antagonist control** | testolactone | 13769 | C_19_H_24_O_3_ | Yes  -97.4206 |  |  |  |  |  |
| **17. Estrogen receptor alpha antagonist**  **control** | CHEMBL4647229 | 156021126 | C_26_H_32_N_2_O_5_ |  | Yes  -117.612 |  |  |  |  |
| **18. Estrogen receptor alpha antagonist**  **control** | tamoxifen | 2733526 | [C_26_H_29_NO](https://pubchem.ncbi.nlm.nih.gov/#query=C26H29NO) |  | Yes  -111.095 |  |  |  |  |
| **19. HER2 antagonist control** | Lapatinib | 208908 | C_29_H_26_ClFN_4_O_4_S |  |  | Yes  -143.425 |  |  |  |
| **20. HER2 antagonist control** | Neratinib | 9915743 | [C_30_H_29_ClN_6_O_3_](https://pubchem.ncbi.nlm.nih.gov/#query=C30H29ClN6O3) |  |  | Yes  -125.739 |  |  |  |
| **21. HER2 antagonist control** | Dacomitinib | 11511120 | [C_24_H_25_ClFN_5_O_2_](https://pubchem.ncbi.nlm.nih.gov/#query=C24H25ClFN5O2) |  |  | Yes  -114.611 |  |  |  |
| **22. MT2 agonist control** | Ramelteon | 208902 | [C_16_H_21_NO_2_](https://pubchem.ncbi.nlm.nih.gov/#query=C16H21NO2) |  |  |  | Yes  -112.932 |  |  |
| **23. PARP10**  **Antagonist**  **control** | Olaparib | 23725625 | C_24_H_23_FN_4_O_3_ |  |  |  |  | Yes  -146.085 |  |
| **24. PARP10**  **Antagonist**  **control** | Talazoparib | 135565082 | C_19_H_14_F_2_N_6_O |  |  |  |  | Yes  -153.637 |  |
| **25. PARP10**  **Antagonist control** | Niraparib | 24958200 | C_19_H_20_N_4_O |  |  |  |  | Yes  -107.072 |  |
| **26. STING agonist**  **control** | STING-agonist-C11 | 16272610 | [C_19_H_18_N_4_O_3_S](https://pubchem.ncbi.nlm.nih.gov/#query=C19H18N4O3S) |  |  |  |  |  | Yes  -146.381 |
| **27. STING agonist**  **control** | STING agonist-1 | 5077622 | C_21_H_16_ClFN_2_O_3_S |  |  |  |  |  | Yes  -126.657 |
| **28. STING agonist**  **control** | STING agonist-14 | 156118667 | [C_16_H_15_NO_2_](https://pubchem.ncbi.nlm.nih.gov/#query=C16H15NO2) |  |  |  |  |  | Yes  -106.02 |

**Table 1**
