## Supplementary material for "In Silico Discovery of Multi-Target Natural Ligands and Efficient siRNA Design for Overcoming Drug Resistance in Breast Cancer via Local Therapy": CYP Inhibition Profiles, Mutagenicity, and Cytotoxicity ,Druglikeness

| **Ligand Number** | **CYP1A2 inhibitor** | **CYP2C19 inhibitor** | **CYP2C9 inhibitor** | **CYP2D6 inhibitor** | **CYP3A4 inhibitor** | **CYP2E1 inhibitor** | **Mutagenicity** | **Cytotoxicity** |
| --- | --- | --- | --- | --- | --- | --- | --- | --- |
| **1** | Inactive | Inactive | Inactive | Inactive | Inactive | Inactive | Inactive | Inactive |
| **2** | Inactive | Inactive | Inactive | Inactive | Inactive | Inactive | Inactive | Inactive |
| **3** | Inactive | Active | Active | Inactive | Active | Inactive | Inactive | Inactive |
| **4** | Inactive | Active | Active | Inactive | Inactive | Inactive | Inactive | Inactive |
| **5** | Inactive | Inactive | Inactive | Inactive | Inactive | Inactive | Inactive | Inactive |
| **6** | Inactive | Inactive | Inactive | Inactive | Inactive | Inactive | Inactive | Inactive |
| **7** | Inactive | Inactive | Inactive | Inactive | Inactive | Inactive | Inactive | Inactive |
| **8** | Inactive | Inactive | Inactive | Inactive | Inactive | Inactive | Inactive | Inactive |
| **9** | Inactive | Inactive | Active | Inactive | Inactive | Inactive | Inactive | Inactive |
| **10** | Inactive | Inactive | Inactive | Inactive | Inactive | Inactive | Inactive | Inactive |
| **11** | Inactive | Inactive | Active | Inactive | Inactive | Inactive | Inactive | Inactive |
| **12** | Inactive | Inactive | Active | Inactive | Inactive | Inactive | Inactive | Inactive |
| **13** | Inactive | Inactive | Active | Inactive | Inactive | Inactive | Inactive | Inactive |
| **14** | Active | Inactive | Inactive | Inactive | Active | Inactive | Inactive | Inactive |
| **15** | Inactive | Inactive | Active | Inactive | Inactive | Inactive | Inactive | Inactive |
| **16** | Inactive | Inactive | Inactive | Inactive | Inactive | Inactive | Inactive | Inactive |
| **17** | Inactive | Inactive | Inactive | Inactive | Inactive | Inactive | Inactive | Inactive |
| **18** | Inactive | Inactive | Active | Inactive | Active | Inactive | Inactive | Inactive |
| **19** | Inactive | Inactive | Active | Active | Active | Inactive | Inactive | Active |
| **20** | Inactive | Inactive | Inactive | Inactive | Inactive | Inactive | Inactive | Inactive |
| **21** | Inactive | Inactive | Active | Active | Inactive | Inactive | Active | Inactive |
| **22** | Inactive | Inactive | Inactive | Active | Inactive | Inactive | Inactive | Inactive |
| **23** | Inactive | Inactive | Active | Inactive | Inactive | Inactive | Inactive | Inactive |
| **24** | Inactive | Inactive | Inactive | Inactive | Inactive | Inactive | Inactive | Inactive |
| **25** | Inactive | Inactive | Inactive | Active | Inactive | Inactive | Inactive | Inactive |
| **26** | Inactive | Inactive | Inactive | Inactive | Inactive | Inactive | Inactive | Inactive |
| **27** | Inactive | Inactive | Active | Inactive | Inactive | Inactive | Inactive | Inactive |
| **28** | Active | Active | Active | Active | Active | Inactive | Active | Inactive |

**Table 2**

| **Ligand Number** | **Number of hydrogen bond donors** | **Number of hydrogen bond acceptors** | **Molweight** | **Lipophilicity (Log Po/w):**  **iLOGP/XLOGP3/WLOGP/MLOGP/SILICOS-IT/Consensus** | **Druglikeness**  **(Lipinski's Rule)** | **Log S (ESOL)/Class** | **GI absorption** | **Predicted LD50** |
| --- | --- | --- | --- | --- | --- | --- | --- | --- |
| **1** | 11 | 18 | 634.45 | 0.98/0.07/-0.30/-2.42/-2.15/.0.76 | No; 3 violations | -3.92/Soluble | Low | 2260mg/kg |
| **2** | 7 | 14 | 593.58 | 2.46/-0.62/-0.64/-1.09/0.48/0.12 | No; 3 violations | -2.38/Soluble | Low | 2000mg/kg |
| **3** | 4 | 9 | 494.45 | 3.23/3.70/3.14/-0.10/3.34/2.66 | Yes; 0 violation | -5.36/Moderately soluble | Low | 5000mg/kg |
| **4** | 3 | 3 | 586.89 | 6.01/12.61/10.28/7.45/9.74/9.23 | No; 2 violations | -10.93/Insoluble | Low | 2510mg/kg |
| **5** | 0 | 2 | 468.75 | 5.44/9.76/8.76/6.98/7.53/7.70 | Yes; 1 violation | -8.43/Poorly soluble | Low | 1185mg/kg |
| **6** | 10 | 12 | 578.52 | 1.49/2.37/2.35/-0.26/1.14/1.42 | No; 3 violations | -5.14/Moderately soluble | Low | 2500mg/kg |
| **7** | 1 | 2 | 416.68 | 5.71/10.33/8.53/5.94/9.20/7.94 | Yes; 1 violation | -8.29/ Poorly soluble | Low | 5000mg/kg |
| **8** | 1 | 2 | 282.46 | 4.27/7.64/6.11/4.57/5.95/5.71 | Yes; 1 violation | -5.41/Moderately soluble | High | 48mg/kg |
| **9** | 1 | 1 | 410.68 | 4.96/7.97/7.72/6.53/6.85/6.81 | Yes; 1 violation | -7.08/Poorly soluble | Low | 10mg/kg |
| **10** | 9 | 12 | 564.49 | 0.66/2.83/1.56/-0.79/1.56/1.07 | No; 3 violations | -5.12/Moderately soluble | Low | 2500mg/kg |
| **11** | 1 | 1 | 412.69 | 4.96/9.45/8.09/6.62/7.18/7.26 | Yes; 1 violation | -7.96/Poorly soluble | Low | 640mg/kg |
| **12** | 1 | 1 | 412.69 | 5.07/9.27/7.94/6.52/7.05/7.19 | Yes; 1 violation | -7.84/Poorly soluble | Low | 890mg/kg |
| **13** | 1 | 1 | 414.71 | 5.03/9.34/8.02/6.73/7.04/7.23 | Yes; 1 violation | -7.90/Poorly soluble | Low | 890mg/kg |
| **14** | 0 | 4 | 285.3 | 2.20/2.73/2.66/1.49/2.53/2.32 | Yes; 0 violation | -3.70/Soluble | High | 1463mg/kg |
| **15** | 0 | 2 | 296.4 | 2.79/3.07/4.03/3.55/4.12/3.51 | Yes; 0 violation | -3.61/Soluble | High | 2920mg/kg |
| **16** | 0 | 3 | 300.39 | 2.64/3.03/3.59/3.34/3.36/3.19 | Yes; 0 violation | -3.61/Soluble | High | 1000mg/kg |
| **17** | 2 | 7 | 452.54 | 4.24/4.82/3.52/2.44/3.95/3.79 | Yes; 0 violation | -5.29 /Moderately soluble | High | 500mg/kg |
| **18** | 0 | 2 | 371.52 | 4.64/7.14/6.00/5.10/5.99/5.77 | Yes; 1 violation | -6.59 /Poorly soluble | Low | 1190mg/kg |
| **19** | 2 | 7 | 581.06 | 4.20/5.12/7.34/3.44/5.84/5.19 | Yes; 1 violation | -6.44 /Poorly soluble | Low | 1500mg/kg |
| **20** | 2 | 9 | 557.04 | 3.96/4.87/5.59/1.64/5.14/4.24 | Yes; 1 violation | -5.98 / Moderately soluble | Low | 4000mg/kg |
| **21** | 2 | 7 | 469.94 | 4.39/4.43/5.00/3.49/3.39/4.34 | Yes; 0 violation | -5.3/Moderately soluble | High | 1000mg/kg |
| **22** | 1 | 3 | 259.34 | 2.90/2.70/2.57/2.49/3.86/2.90 | Yes; 0 violation | -3.05 /Soluble | High | 1000mg/kg |
| **23** | 1 | 6 | 434.46 | 2.84/1.90/1.94/3.09/4.14/2.78 | Yes; 0 violation | -3.70 /Soluble | High | 500mg/kg |
| **24** | 2 | 5 | 380.35 | 2.41/2.25/2.57/3.16/3.12/2.70 | Yes; 0 violation | -4.04 /Moderately soluble | High | 500mg/kg |
| **25** | 2 | 4 | 320.39 | 2.23/2.22/2.21/2.47/2.34/2.29 | Yes; 0 violation | -3.49 /Soluble | High | 2000mg/kg |
| **26** | 2 | 7 | 382.44 | 2.62/3.82/3.01/2.51/2.80/2.95 | Yes; 0 violation | -4.56 /Moderately soluble | High | 4000mg/kg |
| **27** | 1 | 5 | 430.88 | 3.19/3.69/4.38/3.26/4.84/3.87 | Yes; 0 violation | -4.87 /Moderately soluble | High | 1400mg/kg |
| **28** | 1 | 2 | 253.3 | 2.65/3.68/3.31/2.22/4.59/3.29 | Yes; 0 violation | -4.21 /Moderately soluble | High | 3000mg/kg |

**Tabe 3**

| **Plant Name** | **Plant Name** | **Plant Name** | **Plant Name** |
| --- | --- | --- | --- |
| Azadirachta indica L | Thymus pubescens | Achillea santolina | Cymbopogon olivieri |
| Thymus daenensis | Malva sylvestris L. | Juglans regia | Pennyroyal |
| Thymus vulgaris | Licorice (Glycyrrhizaglabra) | Pimpinela anisum L. | Mentha |
| Thymus eriocalyx | Echium italicum L. | Urtica dioica L. | Sage (Salvia officinalis L.) |
| Thymus migricus | Allium jesdianum | Tagetes minuta L. | Mentha longifolia |
| Thymus serpylum | Artemisia absinthium | Ocimum basilicum | Moringaceae |
| Thymus multiflora | Berberis vulgaris | Astragalus calliphysa Bge | Trachyspermum copticum |
| Thymus kotschyanus | Ferula assa-foetida L. | Syzygiumaromaticum | Almond (Prunus amygdalis var. dulcis) |
| Tea | Pistacia atlantica subsp. kurdica | Foeniculum  Vulgare and Cinnamomum Verum | Zingiber officinale |

**Table 4**

| **Ligand Number** | **Source In This Study** | **Ligand Number** | **Source In This Study** |
| --- | --- | --- | --- |
| **1** | Juglans regia | **8** | Moringaceae seeds |
| **2** | Juglans regia | **9** | almond (Prunus amygdalis var. dulcis) |
| **3** | Berberis vulgaris | **10** | Tea |
| **4** | Urtica dioica L | **11** | almond (Prunus amygdalis var. dulcis) |
| **5** | almond (Prunus amygdalis var. dulcis) | **12** | almond (Prunus amygdalis var. dulcis) |
| **6** | Juglans regia | **13** | almond (Prunus amygdalis var. dulcis) |
| **7** | almond (Prunus amygdalis var. dulcis) |  |  |

**Table 5**

| **Receptor** | **PubChem CID of Co-crystal ligand** | **RMSD** |
| --- | --- | --- |
| Aromatase | 53629486 | 0.3245 |
| ERa | 449459 | 0.4238 |
| Mt2 | 445639 | 0.6296 |
| PARP | 11960529 | 0.5261 |
| STING | 168678201 | 0.3201 |
| HER2 | 164575857 | 1.0851 |

**Table 6**
